# D614G mutation alters SARS-CoV-2 spike conformational dynamics and protease cleavage susceptibility at the S1/S2 junction

**DOI:** 10.1101/2020.10.11.335299

**Authors:** Sophie M-C. Gobeil, Katarzyna Janowska, Shana McDowell, Katayoun Mansouri, Robert Parks, Kartik Manne, Victoria Stalls, Megan Kopp, Rory Henderson, Robert J Edwards, Barton F. Haynes, Priyamvada Acharya

## Abstract

The SARS-CoV-2 spike (S) protein is the target of vaccine design efforts to end the COVID-19 pandemic. Despite a low mutation rate, isolates with the D614G substitution in the S protein appeared early during the pandemic, and are now the dominant form worldwide. Here, we analyze the D614G mutation in the context of a soluble S ectodomain construct. Cryo-EM structures, antigenicity and proteolysis experiments suggest altered conformational dynamics resulting in enhanced furin cleavage efficiency of the G614 variant. Furthermore, furin cleavage altered the conformational dynamics of the Receptor Binding Domains (RBD) in the G614 S ectodomain, demonstrating an allosteric effect on the RBD dynamics triggered by changes in the SD2 region, that harbors residue 614 and the furin cleavage site. Our results elucidate SARS-CoV-2 spike conformational dynamics and allostery, and have implications for vaccine design.

**Highlights:**

- SARS-CoV-2 S ectodomains with or without the K986P, V987P mutations have similar structures, antigenicity and stability.
- The D614G mutation alters S protein conformational dynamics.
- D614G enhances protease cleavage susceptibility at the S protein furin cleavage site.
- Cryo-EM structures reveal allosteric effect of changes at the S1/S2 junction on RBD dynamics.

## Introduction

The severe acute respiratory coronavirus 2 (SARS-CoV-2) belongs to the β-coronavirus family of enveloped, positive-sense single stranded RNA viruses, and has one of the largest genomes among RNA viruses (de Wit et al., 2016). Of the seven known coronaviruses that infect humans, four (HCov-229E, HCoV-OC43, HCoV-NL63, CoV-HKU1) circulate annually causing generally mild respiratory symptoms in otherwise healthy individuals, while the SARS-CoV-1 and Middle East respiratory syndrome coronavirus (MERS-CoV), that are closely related to SARS-CoV-2, have resulted in the 2002-2003 SARS and 2012 MERS epidemics (Zumla et al., 2016), respectively. The ongoing pandemic of coronavirus disease of 2019 (COVID-19), is a global public health emergency with more than 37 million cases and 1 million deaths recorded worldwide (Dong et al., 2020) (https://coronavirus.jhu.edu).

The surface of the SARS-CoV-2 is decorated with the spike (S) glycoprotein (Ke et al., 2020; Turonova et al., 2020) that is the target of most current vaccine development efforts (Corbett et al., 2020; Sempowski et al., 2020). In its prefusion conformation the SARS-CoV-2 S protein is a large homo-trimeric glycoprotein forming a crown (from the Latin *corõna*) at the surface of the virus capsid. Each S protomer is subdivided into two domains, S1 and S2, which are delimited by a furin cleavage site at residue 682–685 (Figure 1). The S1 domain comprises the N-terminal domain (NTD), an NTD-to-RBD linker (N2R), the receptor binding domain (RBD), and subdomains 1 and 2 (SD1 and SD2). The S2 domain contains a second protease cleavage site (S2’) followed by the fusion peptide (FP), heptad repeat 1 (HR1), the central helix (CH), the connector domain (CD), heptad repeat 2 (HR2), the transmembrane domain (TM) and a cytoplasmic tail (CT) (Figure 1). The S1 domain is responsible for recognition and binding to the host-cell angiotensin-converting enzyme 2 (ACE2) receptor. The S2 domain is responsible for viral-host-cell membrane fusion and undergoes large conformational changes (Hoffmann et al., 2020a), but only upon furin cleavage and further essential processing by cleavage at the S2’ site by TMPRSS2 and related proteases (Bestle et al., 2020; Hoffmann et al., 2020b; Matsuyama et al., 2020). Previous reports have demonstrated the central role of the dynamics of the RBD domains between a “closed” (or all RBD-down receptor inaccessible conformation) and “open” (or RBD-up) conformations for recognition and binding to the host cell ACE2 receptor (Gui et al., 2017; Shang et al., 2020; Yuan et al., 2017).

**Figure 1.**
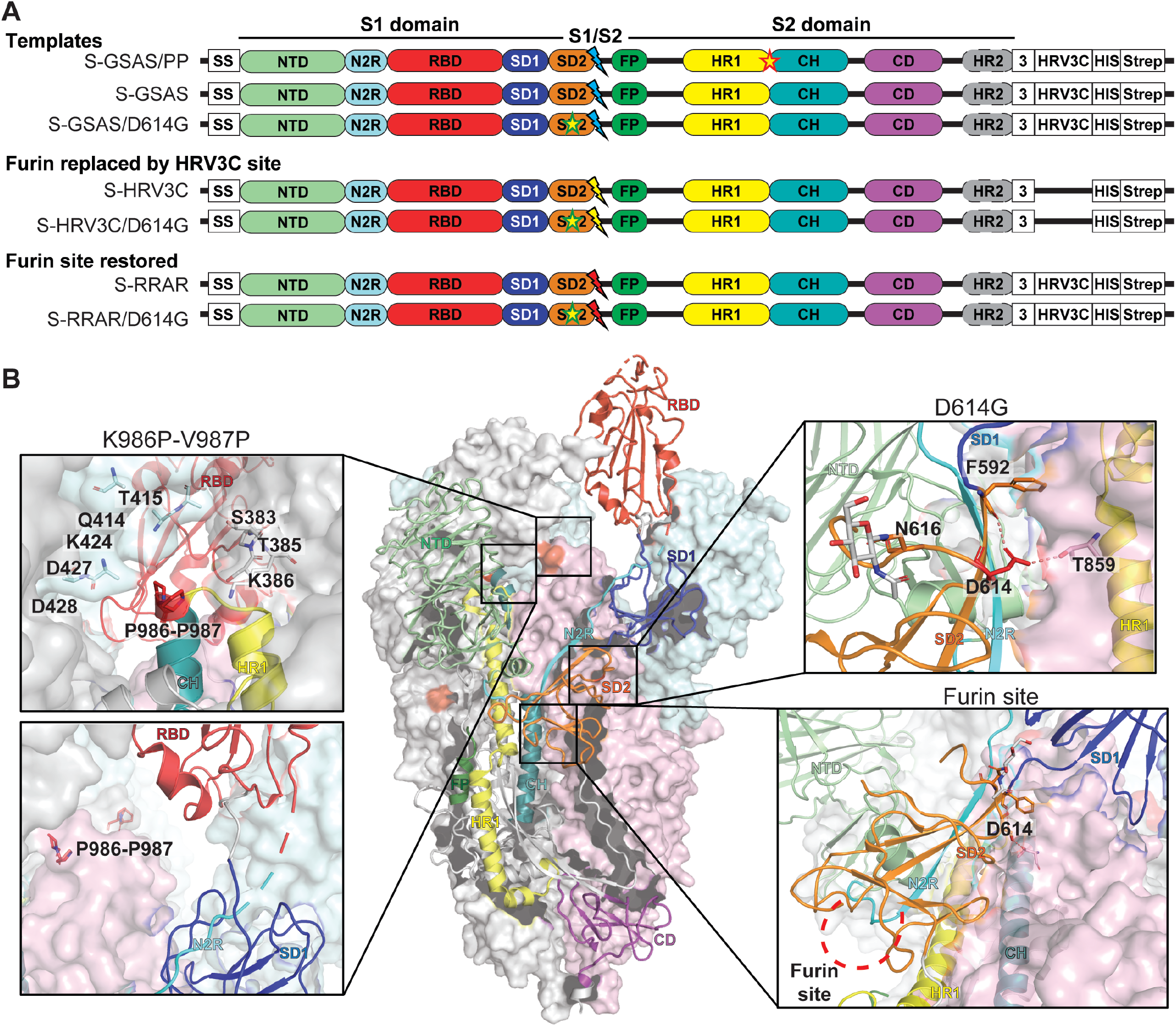
SARS-CoV-2 spike (S) protein ectodomain platform for characterizing the structures, antigenicity and protease susceptibility of the S protein and D614G mutant. **A.** Domain architecture of the SARS-CoV-2 spike protomer. The S1 subunit contains a signal sequence (SS), the NTD (N-terminal domain, pale green), N2R (NTD-to-RBD linker, cyan), RBD (receptor-binding domain, red), SD1 and SD2 (subdomain 1 and 2, dark blue and orange) subdomains. The S2 subunit contains the FP (fusion peptide, dark green), HR1 (heptad repeat 1, yellow), CH (central helix, teal), CD (connector domain, purple) and HR2 (heptad repeat 2, grey) subdomains. The transmembrane domain (TM) and cytoplasmic tail (CT) have been truncated and replaced by a foldon trimerization sequence (3), an HRV3C cleavage site (HRV3C), a his-tag (His) and strep-tag (Strep). The D614G mutation is in the SD2 domain (yellow star, green contour). The S1/S2 furin cleavage site (RRAR; red lightning) has been mutated to GSAS (blue lightning) or to an HRV3C protease cleavage site (yellow lightning). The K986P-V987P mutations between the HR1 and CH domains is indicated by a yellow star (red contour) on the S-GSAS/PP template. **B.** Representation of the trimeric SARS-CoV-2 spike ectodomain with one RBD-up in a prefusion conformation (PDB ID 6VSB). The S1 domain on an RBD-down protomer is shown as pale green molecular surface while the S2 domain is shown in pale red. The subdomains on an RBD-up protomer are colored according to panel **A** on a ribbon diagram. Each inset correspond to the spike regions understudy and are highlighted in red on the trimeric structure (K986P-V987P, D614G and the furin protease cleavage site).

Since the early stages of the COVID-19 pandemic, virus evolution has been followed by large-scale sequencing of the virus genomes isolated from patients, and several mutations that arose and propagated within different populations have been identified even though the virus has genetic proofreading mechanisms (Elbe and Buckland-Merrett, 2017; Korber et al., 2020). The D614G mutation in particular has attracted attention since it has quickly become the dominant strain of SARS-CoV-2 circulating worldwide (Korber et al., 2020). The D614G mutation of the S protein has been associated in numerous reports with increased fitness and/or infectivity of the virus (Korber et al., 2020; Li et al., 2020; Weissman et al., 2020). Cryo electron microscopy (cryo-EM) structures of the S glycoprotein ectodomain have revealed that D614 is a surface residue in the vicinity of the furin cleavage site. Mutation of this residue to a glycine is expected to disrupt a critical interprotomer hydrogen bond involving T859 of the S2 domain (Korber et al., 2020) and resulting in a shift in the observed equilibrium between the open and closed state of the S protein ectodomain (Johnson et al., 2020; Weissman et al., 2020; Yurkovetskiy et al., 2020) (Figure 1).

Most structures of the SARS-CoV-2 S ectodomain currently available include two mutations, one to disrupt the furin cleavage site (RRAR to GSAS = S-GSAS), and a double proline mutation (PP) of residues 986–987, designed to prevent conformational change to the post-fusion state (Wrapp et al., 2020). Originally designed for the MERS S protein (Pallesen et al., 2017), insertion of two consecutive Pro mutations at the junction of the HR1 and CH regions stabilized the pre-fusion conformation of the MERS, SARS and HCoV-HKU spikes, increased protein expression, and immunogenicity for the MERS S protein (Pallesen et al., 2017). Based on these prior data, introduction of two consecutive proline residues at the beginning of the central helix was postulated as a general strategy for retaining β-coronavirus S proteins in the prefusion conformation. Thus, the PP mutations were carried over to the SARS-CoV-2 ectodomain (Wrapp et al., 2020) that is currently widely used in the field for vaccine and structural studies, and is also the component of a vaccine candidate (Corbett et al., 2020). Although shown to stabilize the pre-fusion conformation of other coronaviruses, the effect of the PP insertion has not been systematically studied for the SARS-CoV-2 S ectodomain.

With the goal of investigating the biophysical and structural consequences of the D614G mutation, and to prevent the engineered PP mutations from confounding our observations, we produced two SARS-CoV-2 S ectodomain constructs with the native K986 and V987 residues, incorporating either a D or a G at position 614 (Figure 1). The RRAR sequence in the furin cleavage site was replaced by a GSAS sequence thus rendering the S constructs furin-cleavage deficient. To probe the effect of the D614G substitution on furin cleavage of the S protein, we either reinstated the native furin sequence or replaced it with an exogeneous HRV3C proteolysis signal. We determined the cryo-EM structures of the uncleaved D614 and G614 S ectodomains, as well as the structure of the fully cleaved G614 S ectodomain of the currently globally dominant SARS-CoV-2. Our results demonstrate the effect of the D614G substitution on the conformational dynamics and furin cleavage susceptibility of the S ectodomain, and reveal insights into the allostery between the RBD and distal regions of the S protein.

## Results

### Structure and stability of the SARS-CoV-2 S ectodomain incorporating the native K986 and V987 residues

While the SARS-CoV-2 S ectodomain construct that includes mutations of residues K986 and V987, between the HR1 and CH subdomains (S2 domain), to prolines (PP) (named S-GSAS/PP in this study) (Figure 1) is widely used in the field, the origin of this PP construct was based upon the stabilization of the pre-fusion conformation of other coronavirus spikes (Pallesen et al., 2017; Walls et al., 2020; Wrapp et al., 2020). Here, we generated an analogous S ectodomain construct that had the native K986 and V987 residues (named S-GSAS) (Figure 1).

In our 293F expression system (see methods for details), both the S-GSAS/PP and S-GSAS constructs expressed at similar levels, yielding about 3 mg final protein per L of culture. Both proteins also showed similar migration profiles on SDS-PAGE and by size exclusion chromatography (SEC) on a Superose 6 column (Figure 2A, B).

**Figure 2.**
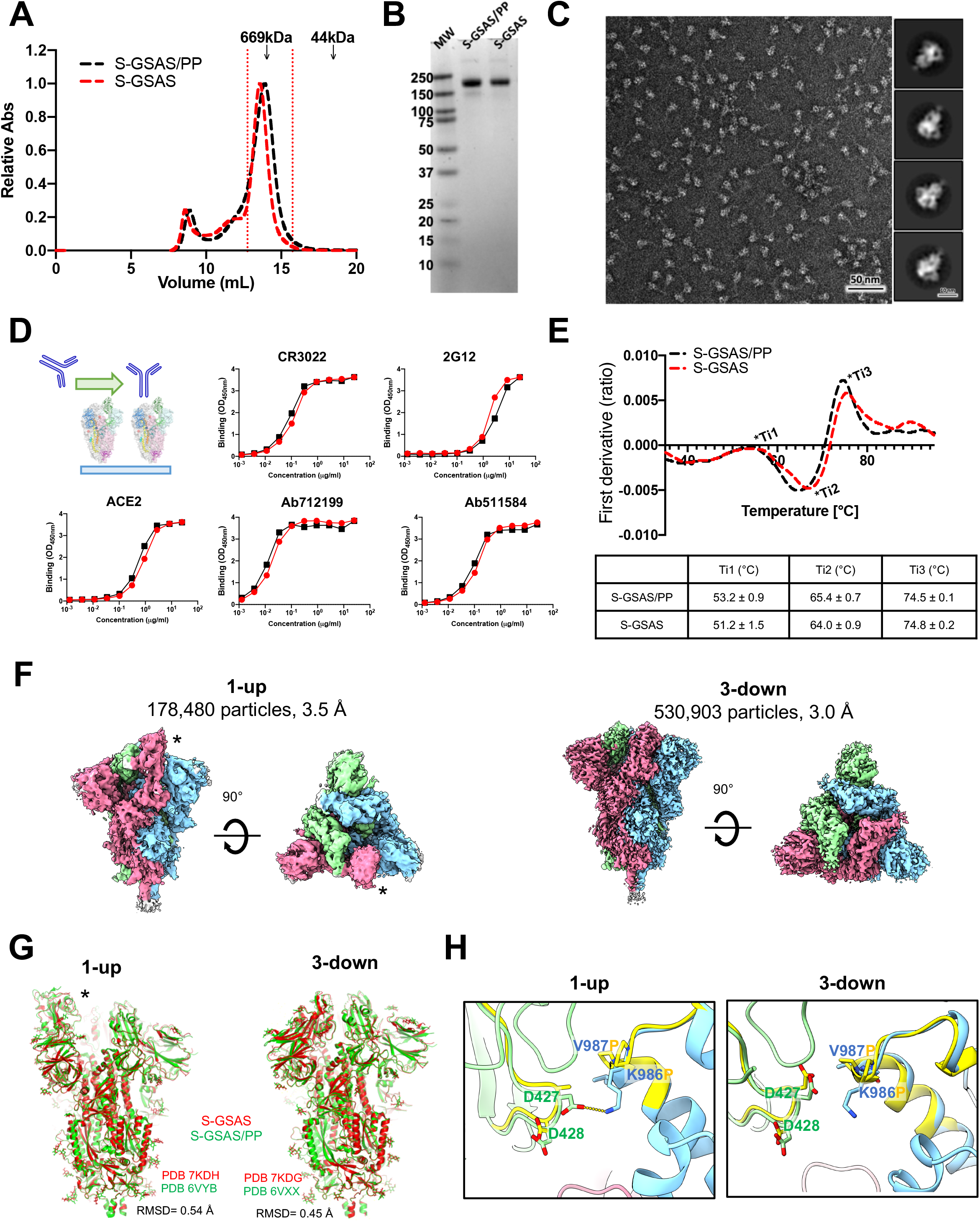
Biophysics, antigenicity and structure of the S-GSAS ectodomain in relation to S-GSAS/PP. **A.** Size-exclusion chromatography (SEC) elution profile on a superose 6 10/300 column of the S-GSAS/PP (black) and S-GSAS (red) ectodomains. Fractions isolated for further characterization are indicated by vertical red dotted lines. Elution volumes of molecular weight standards at 669 (thyroglobulin) and 44 kDa (ovalbumin) are labelled for reference. **B.** SDS-PAGE of the SEC purified ectodomains. **C.** Representative NSEM micrograph of S-GSAS and 2D class averages. **D.** Binding of ACE2 receptor ectodomain (RBD-directed), CR3022 (RBD-directed neutralizing antibody), 2G12 (S2-directed), Ab712199 (RBD-directed neutralizing antibody) and Ab511584 (S2-directed non-neutralizing antibody) to S-GSAS (red) and S-GSAS/PP (black) measured by ELISA. The schematic shows the assay format. Serially diluted spike protein was bound in individual wells of 384-well plates, which were previously coated with streptavidin. Proteins were incubated and washed, then antibodies at 10 μg/ml or ACE2 with a mouse Fc tag at 2 μg/ml were added. Antibodies were incubated, washed and binding detected with goat anti-human-HRP. **E.** Differential scanning fluorimetry (DSF) of the S-GSAS (red) and S-GSAS/PP (black) S ectodomains. Thermal melting inflection points (Ti) are indicated on the first derivative graph and reported in the table below. **F.** Side and top view of the cryo-EM reconstructions of the 1-RBD-up and the 3-RBD-down states of the S-GSAS ectodomain colored by chain. The up positioned RBD in the map is identified by an asterisk. **G.** Superposition of the 1-up (left) and 3-down (right) structures of S-GSAS (red) and S-GSAS/PP (green). **H.** Magnified view of one protomer from the 1-RBD up model showing residues K986 and V987 from S-GSAS (colored according to panel **F**, overlayed with S-GSAS/PP (PDB 6VYB; yellow)), showing residues P986 and P987 in sticks.

Similar to the S-GSAS/PP construct (Edwards et al., 2020; Henderson et al., 2020; Wrapp et al., 2020), S-GSAS showed 80–90% intact prefusion spike trimers by negative stain electron microscopy (NSEM) (Figure 2C). This finding is in contrast to previous observations for MERS and the SARS-CoV-1 ectodomains, which showed a mixture of the prefusion and postfusion conformations unless the PP mutation was included (Pallesen et al., 2017). Binding of S-GSAS and S-GSAS/PP measured by ELISA to ACE2 and CR3022 (Edwards et al., 2020), both requiring an RBD-up conformation, Ab712199 and Ab511584, two antibodies isolated from a COVID-19 convalescent donor with epitopes mapping to the ACE2 binding site and S2 domain respectively, and 2G12, binding to a quaternary S2 glycan epitope, were all nearly identical demonstrating that both constructs showed similar antigenic behavior (Figure 2D). Using differential scanning fluorimetry to measure the spike thermostability, we found the S-GSAS and S-GSAS/PP ectodomains showed similar melting temperatures (Figure 2E).

We next solved cryo-EM structures of the S-GSAS ectodomain (Figures 2F-H, **S1-S2, Table S1**), to compare with the S-GSAS/PP structures (Walls et al., 2020; Wrapp et al., 2020) and to visualize the impact that the engineered PP mutations had on the structure of the SARS-CoV-2 spike ectodomain. Two populations of the S-GSAS S ectodomain were identified in the cryo-EM dataset - a 1-RBD-up (or open) and a 3-RBD-down (or closed) conformation (Figure 2F and **Table S1**). Both structures were similar to the corresponding structures of S-GSAS/PP (Walls et al., 2020), with overall RMSDs of 0.45 Å and 0.54 Å for the 1-up and 3-down structures, respectively. In the region around the PP mutations, we found the S-GSAS structures to be similar to the corresponding S-GSAS/PP structures (Figure 2H). In the S-GSAS 1-RBD-up structure, we observed that the K986 side chain was appropriately positioned to make an interprotomer salt bridge with the D427 residue of the RBD of the adjacent protomer, an interaction that would be abrogated in the PP construct. The corresponding residues in the MERS S protein, V1060 and L1061 are non-polar, and the adjacent protomers too far to interact with these residues (**Figure S3**). In the SARS-CoV-1 S protein cryo-EM structure (PDB 5XLR, 5X5B) the residues D414-D415 (equivalent to SARS-CoV-2 D427-D428) lie further from K986 suggesting that this putative salt bridge interaction may be more transient in SARS-CoV-1.

Overall, our data show that for the SARS-CoV-2 S ectodomain, the S-GSAS construct showed similar structural, antigenic and stability behavior as the S-GSASS/PP construct that included the K986P and V987P mutations at the junction of the CH and HR1 regions. While these and analogous mutations had proved beneficial for the expression and stability of other CoVs (Pallesen et al., 2017), for the SARS-CoV-2 S protein other compensating interactions may help confer stability to the pre-fusion form in the absence of the PP mutations. For the rest of this study we have used the S-GSAS construct as the platform for introducing mutations and other modifications of interest.

### The SARS-CoV-2 Spike glycoprotein D614G mutation

To understand the molecular details of the spike D614G mutation that arose and quickly dominated circulating SARS-CoV-2 isolates globally, we sought to assess the impact of the D614G mutation on the structure and antigenicity of the SARS-CoV-2 S ectodomain. The D614G mutated S-GSAS construct (S-GSAS/D614G), yielded an average of ∼ 2 mg of purified protein per L of culture (n = 4). The SDS-PAGE, SEC and DSF profiles of the S-GSAS/D614G (Figure 3A) were similar to that of the S-GSAS S ectodomain (Figure 2A, B). NSEM of the S-GSAS/D614G S ectodomain revealed typical and well-dispersed pre-fusion S particles (Figure 3B).

**Figure 3.**
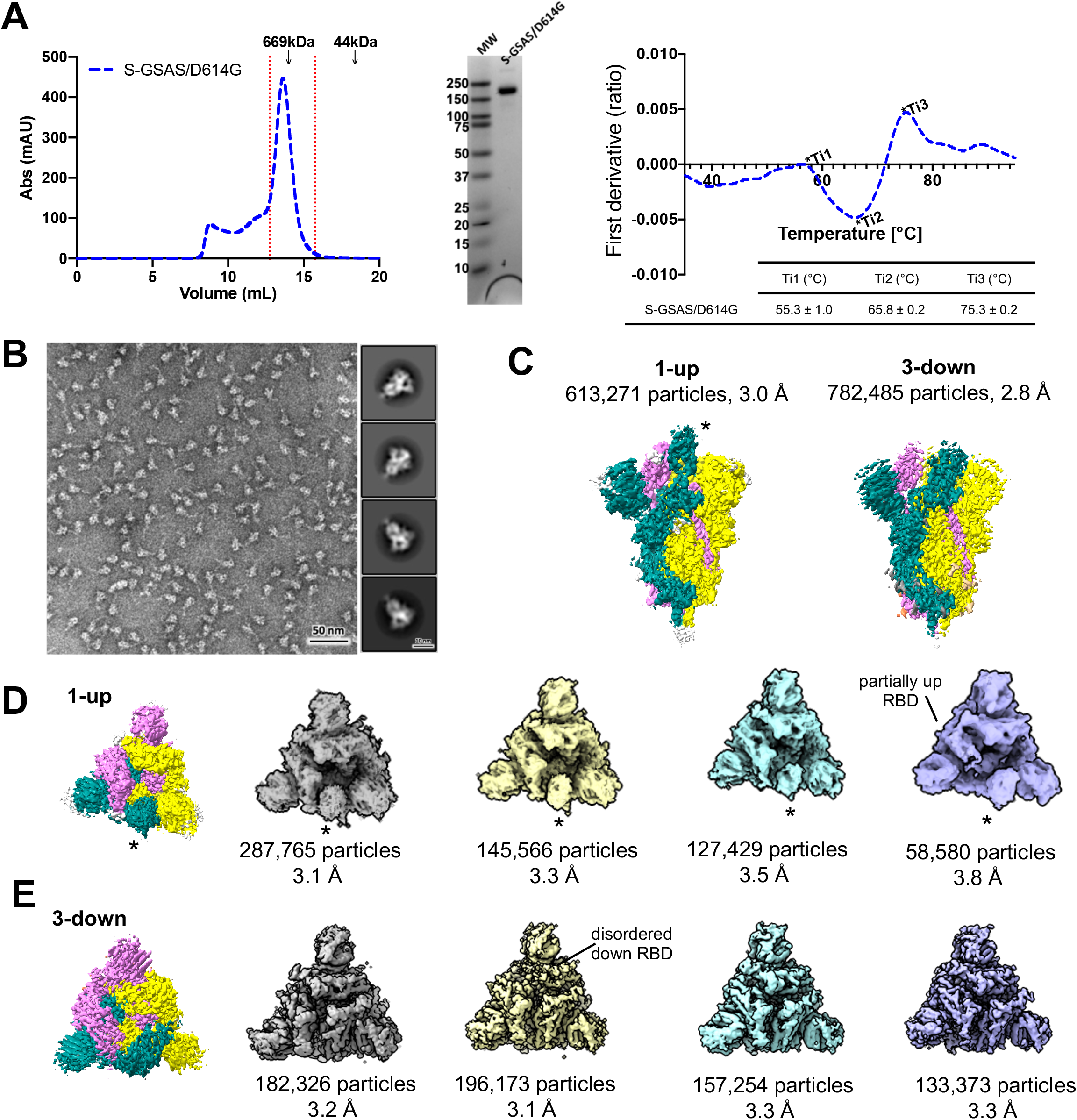
Biophysics and structure of the S-GSAS/D614G ectodomain. **A.** (left) Size-exclusion chromatography (SEC) elution profile on a superose 6 10/300 column of the S-GSAS/D614G (blue) ectodomain. Fractions isolated for further characterization are indicated by vertical red dotted lines. Elution volumes of standard at 669 and 44 kDa are labelled for reference. (middle) SDS-PAGE of the SEC purified ectodomain. (right) Differential scanning fluorimetry (DSF) of S-GSAS/D614G (blue). Thermal melting inflection points (Ti) are indicated on the first derivative graph and reported in the table below. **B.** Representative NSEM micrograph of S-GSAS/D614G and 2D class averages. **C**. Side view of the cryo-EM reconstruction of the 1-RBD-up and the 3-RBD-down states of the S-GSAS/D614G ectodomain colored by chain. The up positioned RBD in the map is identified by an asterisk. **D.** (left) top view of the 1-RBD-up S trimer shown in **C**. and (right) subpopulations obtained by further classification. **E.** (left) top view of the 3-RBD-down S trimer shown in **C.** and (right) subpopulations obtained by further classification.

To visualize structural details at higher resolution, we determined the cryo-EM structures of S-GSAS/D614G construct (Figure 3C-E, **Table S1, Figures S4-S7**). Two major populations of the S ectodomain were identified in the cryo-EM dataset - one population with one RBD in the “up” or ACE2 receptor accessible conformation and the other with all three RBDs in the “down” or receptor inaccessible conformation. Despite extensive classifications, including searching with low-pass filtered maps of 2-up and 3-up S ectodomains we could not identify any populations with 2-RBDs or 3-RBDs in the “up” state. This is in contrast with cryo-EM results of the D614G mutation published in the context of a S-GSAS/PP spike (Yurkovetskiy et al., 2020) where 2-RBD-up and 3-RBD-up spike populations were identified. The 1-RBD-up population consisted of 613,271 particles and resolved to an overall resolution of 3.0 Å, while the 3-RBD-down population consisted of 782,485 particles and was refined to an overall resolution of 2.8 Å with C3 symmetry applied (Figure 3C, **Table S1, Figures S4-S5**). Both the up and down populations showed considerable heterogeneity in the S1 subunit, primarily originating from the variability in the positions of the RBD and NTD, which could be partially resolved by further classification and separation of subpopulations with different dispositions of the RBD and NTD even though falling broadly under the 1-up and 3-down categories (Figure 3D-E, **Figures S6-S7**). Spike populations that refined to a global resolution of 3.0 Å or better (Figure 3D-E) were used for model fitting and further analyses. Comparing with the S-GSAS dataset (Figure 2F), we observed an increased proportion of the 1-RBD-up form versus the 3-RBD-down form in the S-GSAS/D614G data (Figure 3C). This is consistent with our previous observations made with NSEM data that showed an increase in the RBD-up population for the S-GSAS/D614G S ectodomain (Weissman et al., 2020). Our results show that the D614G mutation in the SD2 domain, even though distal from the RBD region, has an allosteric effect on RBD dynamics leading to alteration of up/down RBD dispositions.

To understand the nature of this allostery, we examined changes in the S protein that accompany the up and down RBD transition (Figure 4) by comparing the RBD-up chain in the 1-RBD-up structure to the down chains in the 1-up and the 3-down structures (Figure 3C). In each S protein protomer, the polypeptide chain folds into domains as it traverses the length of the S1 subunit and enters the S2 subunit *i.e.* the NTD (residues 27–305) followed by the RBD (residues 335–521), the SD1 (residues 529–591) and SD2 (residues 592–697) domains (Figure 4A). The NTD and RBD are connected via a 28-residue linker spanning residues 306–334 (named N2R) that stacks against the SD1 and SD2 domains (Figure 4A-D), as it makes its way from the NTD to the RBD, essentially connecting all the individual domains in the S1 subunit, and forming “super” subdomains SD1’ and SD2’, respectively (Henderson et al., 2020). Upon overlaying the protomers with the RBD in the up position with the protomers with their RBDs in the down position by using the S2 subunit residues 908–1035 for superpositions, we found that the down-to-up RBD motion is accompanied by a rigid body movement of the SD1’ domain resulting in a shift of up to ∼4.5 Å of the SD1 domain (Figure 4D), relative to its position in the RBD-down protomers and a shift of up to a 7 Å in the N2R linker as it hinges to enter into the RBD. This results in a ∼20° tilt of residues 324–328 of the N2R linker region that forms part of the SD1’ super subdomain, while residues 311–319 of the linker that associate with the SD2 subdomain remained virtually unmoved, showing only a slight tilt in the β beta strand that accompanied large movements in the RBD and adjoining SD1’ domain (Figure 4D). Indeed, the SD2’ super subdomain that harbors the D614G mutation appears to form a conformationally invariant anchor with the mobile RBD and NTD domains at either end (Figure 4D). Additionally, the S2 subunit remains invariant between the different protomers showing that the large movements that occur in the S1 subunit are effectively arrested by the SD2’ super subdomain conformationally invariant anchor.

**Figure 4.**
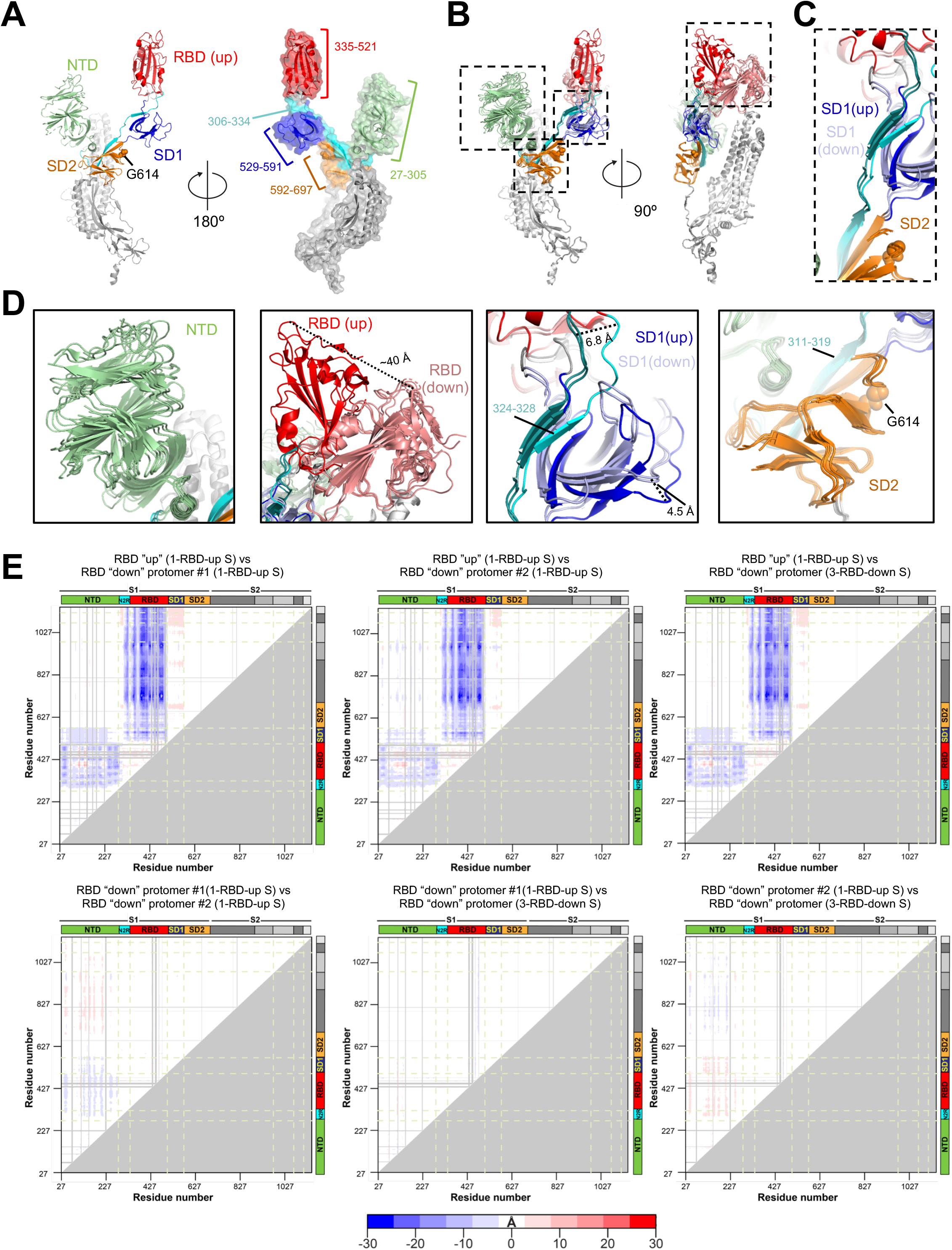
Domain motions in the S-GSAS/D614G ectodomain. **A.** RBD-up chain from the structure shown in Figure 3C with the S1 subunit colored by domain and the S2 subunit colored grey. RBD is colored red, NTD green, SD1 dark blue, SD2 orange and the linker between the NTD and RBD colored cyan. **B.** Overlay of the individual protomers in the 1-RBD-up structure and a protomer in the C3 symmetric 3-RBD-down structure shown in Figure 3C. The structures were superimposed using S2 subunit residues 908–1035 (spanning the HR1 and CH regions). The domain colors of the up-RBD chain are as described in panel **A.** The down-RBDs are colored salmon, the SD1 domains from the down RBD chains are colored light blue. The linker between the NTD and RBD in the down RBD chains are colored deep teal. **C.** Zoomed-in view showing the association of the linker connecting the NTD and RBD with the SD1 and SD2 domains. **D.** Zoomed-in views of individual domains marked in panel **B.** The N2R linker spanning residues 306–334 connects the NTD and the RBD. Residues 324-328 of the N2R linker contribute a β-strand to the SD1 subdomain together forming the SD1’“super” subdomain. Residues 311–319 of the N2R linker contribute a β-strand to the SD2 subdomain together forming the SD2’“super” subdomain. **E.** Difference distance matrices (DDM) showing structural changes between different protomers for the structures shown in Figure 3C. The blue to white to red coloring scheme is illustrated at the bottom.

These observations are mirrored in the difference distance matrices (DDM) comparing the RBD-up and down chains (Figure 4E **and Figure S8**). DDM analyses (Richards and Kundrot, 1988) provide superposition-free comparisons between a pair of structures by calculating the differences between the distances of each pair of Cα atoms in a structure and the corresponding pair of Cα atoms in the second structure. The DDM analysis not only shows the large movement in the RBD region and the movement in the NTD, it also captures the movement in the N2R linker and the SD1 domain observed in the structures. Overall, these analyses show that the D614G mutation is acquired within a key region encompassing the SD2 domain and an additional β-strand contributed by residues 311–319 of the N2R linker that forms a region of relative structural stillness separating the mobile NTD and RBD, as well isolating the motions in S1 from the S2 subunit. This distal mutation altering RBD conformational dynamics shows that small changes in this region can translate into large allosteric effects, and suggests a role for the SD2 domain in modulating RBD dynamics.

### Effect of the D614G substitution on furin cleavage efficiency at the S1/S2 junction

In addition to the D614G mutation, the SD2 subdomain also harbors a furin cleavage site (residues 682–685) that separates the S1 and S2 subunits (Figure 1). Cleavage of the S protein by furin at this site is essential for virus transmission (Shang et al., 2020). The proximity of the D614G mutation to the furin cleavage site and the increased flexibility observed in the cryo-EM dataset of the S-GSAS/D614G ectodomain (Figure 3C-E), prompted us to examine the effect of the D614G substitution on furin cleavage.

Since our expression system *(i.e.* 293Freestyle cells) endogenously expresses furin, in order to obtain uncleaved spike that we could then test for protease cleavage *in vitro*, we engineered a HRV3C site (8 amino acids long) to replace the furin cleavage site (4 amino acids long) at the S1/S2 junction, resulting in the S-HRV3C and S-HRV3C/D614G S ectodomain constructs (Figure 1A). Both proteins expressed in 293F cells but at lower yields compared to the S-GSAS constructs (36 μg/L and 410 μg/L for the S-HRV3C and S-HRV3C/D614G proteins, respectively). SEC and SDS-PAGE profiles were similar to the S-GSAS and S-GSAS/D614G proteins confirming well-folded and homogeneous spike preparations (Figure 5A, B). NSEM micrographs showed characteristic kite-shaped particles (Edwards et al., 2020) for the pre-fusion S protein, and 2D-classification of particles from NSEM revealed well folded spikes, further confirming that S-HRV3C spikes retained the overall fold and structure of the S-GSAS spikes (Figure 5C, D).

**Figure 5.**
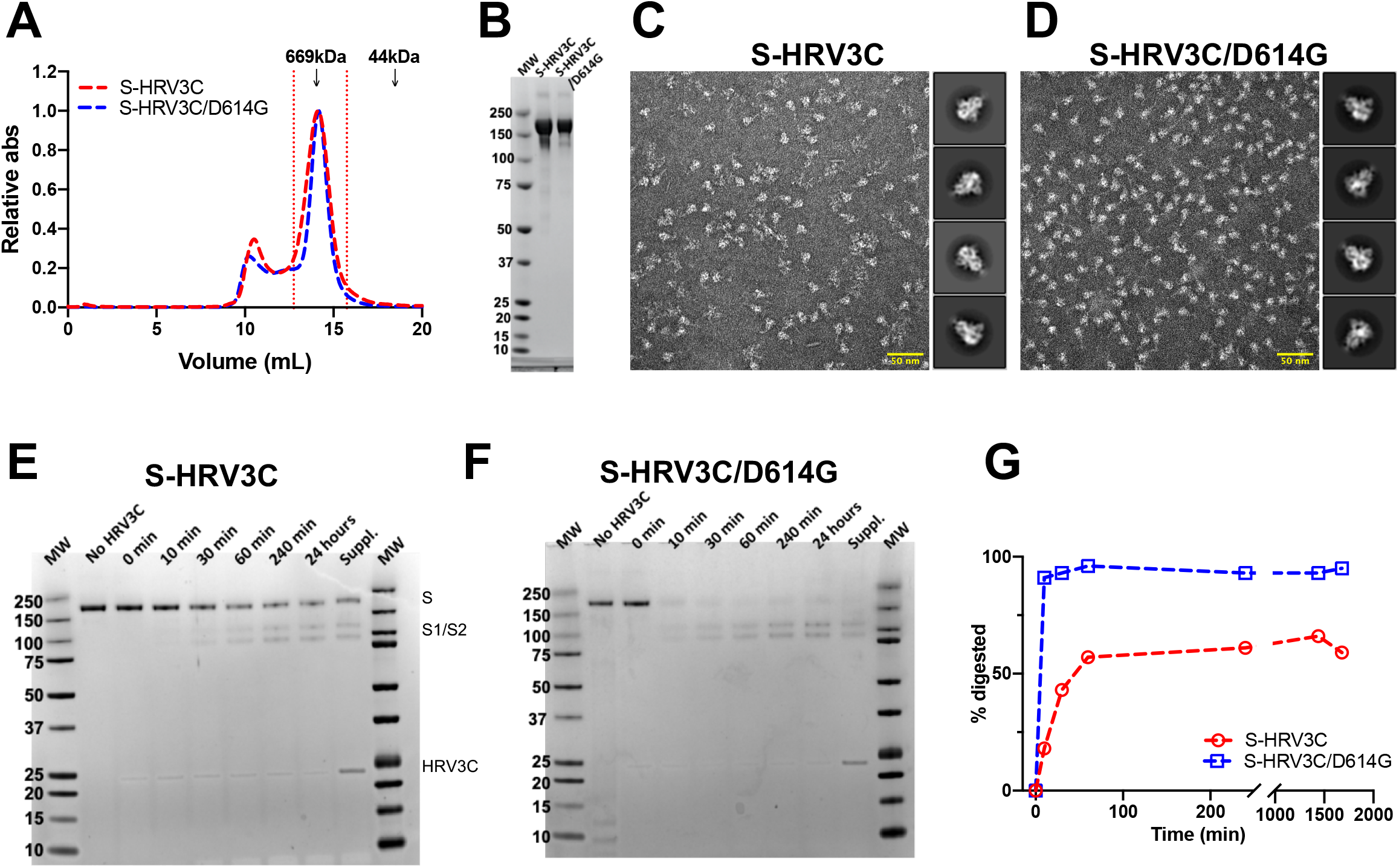
The engineered S-HRV3C/D614G ectodomain is more susceptible to S1/S2 cleavage by the HRV3C protease than S-HRV3C. **A.** Size-exclusion chromatography (SEC) elution profile on a superose 6 10/300 column of the S-HRV3C (red) and S-HRV3C/D614G (blue) ectodomains. Fractions isolated for further characterization are indicated by vertical red dotted lines. Elution volumes of standards at 669 and 44 kDa are labelled for reference. **B.** SDS-PAGE of the SEC purified ectodomains. **C-D.** Representative NSEM micrograph of **C.** S-HRV3C and **D.** S-HRV3C/D614G ectodomains and 2D class averages. **E-F.** SDS-PAGE of an HRV3C digestion of the **E.** S-HRV3C and **F.** S-HRV3C/D614G engineered ectodomains at 25°C for 24 hours in presence of of 0.03 units of enzyme per μg of ectodomain. Aliquots corresponding to 1μg of protein at the timepoints before HRV3C addition, at addition (0 min) and 10, 30, 60, 240 minutes and 24 hours following HRV3C addition are presented. After 24hours, 0.03 supplementary units of the HRV3C enzyme per μg of ectodomain were added and aliquots were analyzed after 4 additional hours of incubation aiming at completion of the digestion (labeled Suppl.). **G.** Quantification of spike protomer (200 kDa) band intensity on SDS-PAGE at the timepoints presented on panel **E** and **F** (S-HRV3C in red, S-HRV3C/D614G in blue).

To test the susceptibility of the HRV3C site engineered at the junction of the S1 and S2 subunits to protease cleavage, we incubated the purified S-HRV3C and S-HRV3C/D614G spikes with the HRV3C enzyme and followed the digestion by analyzing aliquots taken at different time-points by SDS-PAGE (Figure 5E-G). We found that the digestion of the S-HRV3C/D614G spike (Figure 5F-G) proceeded at a faster rate than that of the S-HRV3C spike (Figure 5E-G) with the S-HRV3C/D614G spike almost 100% digested within the first 10 minutes of incubation, whereas, the S-HRV3C constructs only achieved 50% of cleavage after 24 hours, and a substantial portion remained uncleaved even upon addition of more enzyme followed by 4 additional hours of incubation. These results suggested that the D614G mutation increased the susceptibility of protease cleavage at the S1/S2 junction.

To study the effect of the D614G substitution on protease cleavage at the S1/S2 junction with the native furin site, we next generated spike ectodomains constructs where the furin site was restored to the native sequence, resulting in two constructs named S-RRAR and S-RRAR/D614G (Figure 1A). The proteins were expressed and purified using our usual methodology for the furin cleavage-deficient constructs (see methods). The SEC profiles (Figure 6A) showed a higher proportion of the first higher molecular weight peak. A second peak eluting at a similar molecular weight as the S-GSAS spike (at ∼13.8 mL elution volume) was used for further characterization. The SEC profile of the S-RRAR spike preparation showed small populations of lower molecular weight peaks that were not observed for the S-RRAR/D614G protein (Figure 6A). On SDS-PAGE (Figure 6B), the peak corresponding to the S ectodomain showed the S-RRAR construct as having one major band at the molecular weight corresponding to the spike monomer and some fainter bands corresponding to the S1 and S2 subunits while the S-RRAR/D614G protein showed a band corresponding to the spike monomer and the two bands corresponding to the molecular weights of the S1 and S2 subunits. The smaller molecular weight bands corresponding to the S1 and S2 subunits were in higher proportions in the S-RRAR/D614G spike preparation compared to the S-RRAR preparation. In summary the SEC and SDS-PAGE profiles showed that, although both the S-RRAR and S-RRAR/D614G constructs were cleaved by endogeneous furin (Figure 6B) during protein expression the S1 and S2 subunits remained together in solution (Figure 6A). Consistent with the enhanced cleavage observed for the S-HRV3C/D614G spike relative to the S-HRV3C spike, in the furin-site restored spikes we observed a higher proportion of cleaved spike in S-RRAR/D614G relative to S-RRAR, suggesting that the D614G mutation makes the spike more susceptible to furin cleavage. NSEM of the purified S-RRAR (Figure 6C) and S-RRAR/D614G (Figure 6D) confirmed that both of these furin site-restored spikes formed well-folded spike ectodomains.

**Figure 6.**
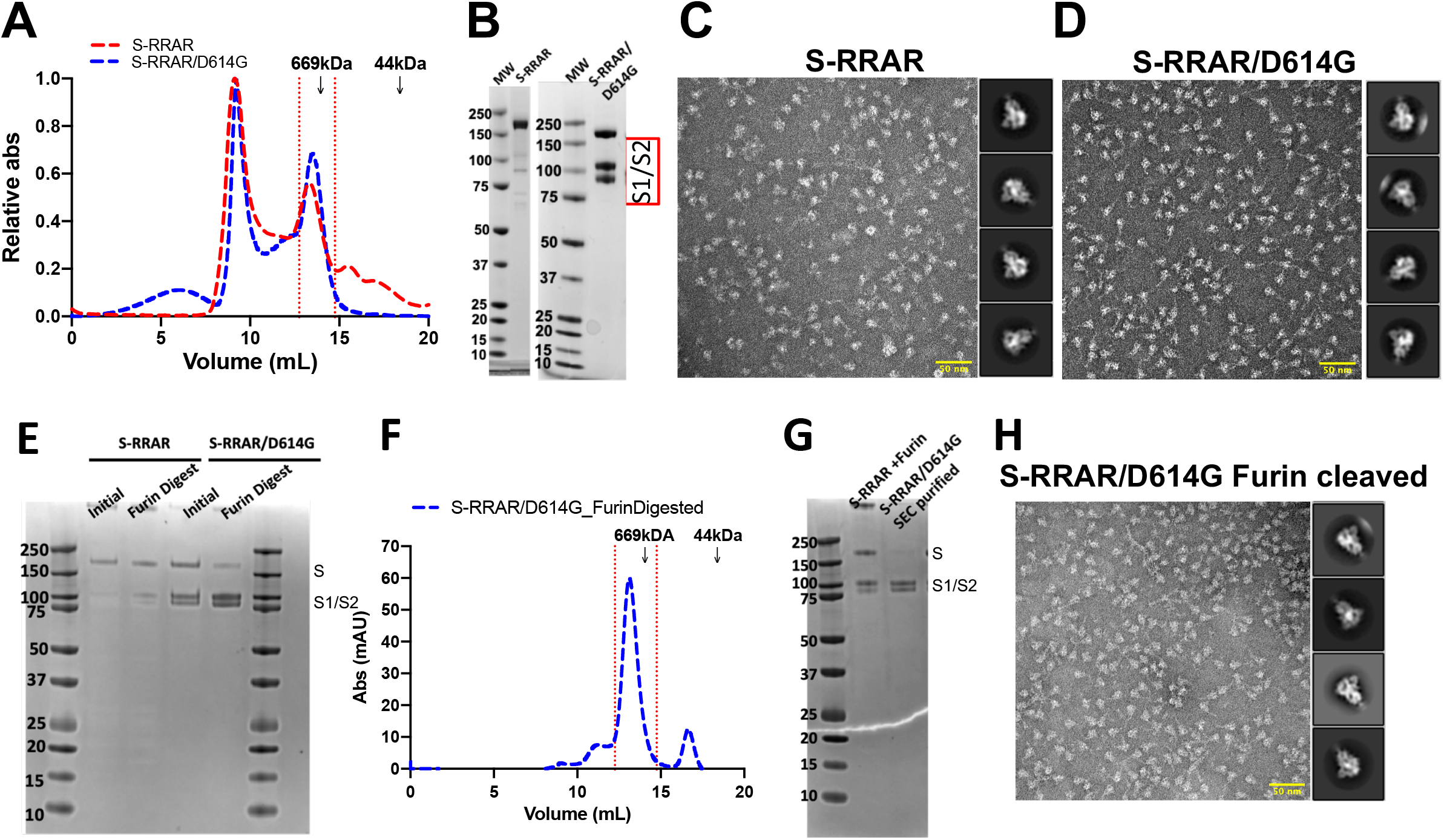
The S-RRAR/D614G ectodomain is more susceptible to S1/S2 cleavage by furin than S-RRAR. **A.** Size-exclusion chromatography (SEC) elution profile of the S-RRAR (in red) and S-RRAR/D614G (in blue) ectodomains. Fractions isolated for further characterization are indicated by vertical red dotted lines. Elution volumes of standards at 669 and 44 kDa are labelled for reference. **B.** SDS-PAGE of the SEC purified ectodomains. The S1 and S2 domains corresponding bands are identified. **C-D.** Representative NSEM micrograph of **C.** S-RRAR and **D.** S-RRAR/D614G ectodomains and 2D class averages. **E.** SDS-PAGE of furin digestion of the S-RRAR and S-RRAR/D614G ectodomains at 25°C for 3 hours in presence of 0.3 units of enzyme per μg of ectodomain in buffer containing 0.2 mM CaCl_2_. Aliquots corresponding to 1μg of protein at the timepoints before furin addition and 3 hours post-addition are presented. **F.** Size-exclusion chromatography (SEC) elution profile of the S-RRAR/D614G furin digested (in blue). Fractions isolated for further characterization are indicated by vertical red dotted lines. **G.** SDS-PAGE of the S-RRAR/D614G furin digested and SEC purified ectodomain. The S1 and S2 domains corresponding bands are identified. In lane 2, the S-RRAR ectodomain was further incubated for 16 hours with 0.3 units of furin per μg of ectodomain aiming at completing the digestion. **H.** Representative NSEM micrograph and 2D class averages of the S-RRAR/D614G furin digested and SEC purified following digestion.

We next digested the SEC isolated fractions of the S-RRAR and S-RRAR/D614G ectodomains (Figure 6A-D) *in vitro* by adding furin (Figure 6E). As observed for the S-HRV3C constructs, the D614 version of the spike was less susceptible to cleavage than the G614 mutant for the same incubation time with the enzyme. SEC purification of the fully digested S-RRAR/D614G ectodomain revealed a peak corresponding to the ectodomain (Figure 6F). On SDS-PAGE, this peak migrated as two distinct bands corresponding to the S1 and S2 domains thus confirming isolation of only the cleaved portion of the protein (Figure 6G). NSEM showed fully folded ectodomains for the furin digested and SEC purified S-RRAR/D614G protein (Figure 6H).

In summary, these results show that acquisition of the D614G mutation the S protein SD2 domain resulted in increased furin cleavage of the S ectodomain.

### Structure and antigenicity of the furin-cleaved D614G S ectodomain

To visualize the structure of the furin-cleaved S ectodomain at atomic level resolution, we obtained a cryo-EM dataset, and resolved two populations of the furin-cleaved S ectodomain - a 1-RBD-up and a 3-RBD-down population (Figure 7A, **Figure S9 and S10 and Table S1**). We observed an increased proportion of the 3-RBD-down S compared to the uncleaved D614G S ectodomain, thus reporting a change in the RBD conformational dynamics upon furin cleavage. Consistent with this result, we observed reduced binding to ligands such as ACE-2 and CR3022 that require the RBD to be in the up conformation for binding (Figure 7B). As expected, decrease in binding was also observed with antibody 712199, isolated from a convalescent COVID-19 donor, with an epitope overlapping with the ACE2 binding site. Antibody 2G12 that binds a quaternary glycan epitope in the S2 subunit showed a small decrease in binding with the furin-cleaved S ectodomain, whereas another COVID-19-derived S2 antibody 511584 showed increase in binding with the furin-cleaved S ectodomain.

**Figure 7.**
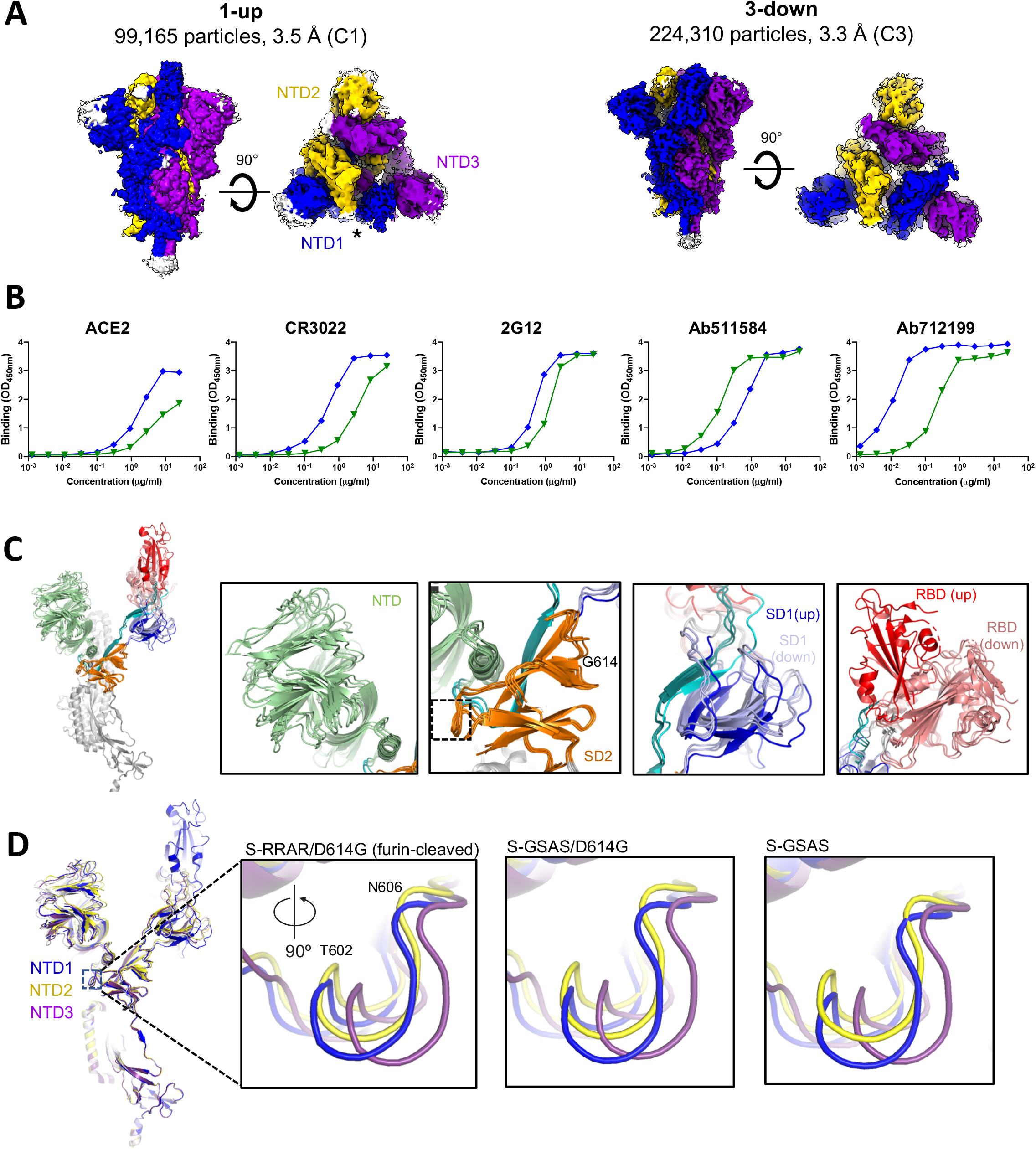
Structure and antigenicity of the furin-cleaved S-RRAR/D614G ectodomain. **A.** Side view of the cryo-EM reconstruction of the 1-RBD-up and the 3-RBD-down states of the furin-cleaved S-RRAR/D614G ectodomain colored by chain. The up positioned RBD in the map is identified by an asterisk. The NTDs in the asymmetric 1-RBD-up structure are labeled. **B.** Binding of ACE2 receptor ectodomain (RBD-directed), CR3022 (RBD-directed neutralizing antibody), 2G12 (S2-directed), Ab712199 (RBD-directed neutralizing antibody) and Ab511584 (S2-directed non-neutralizing antibody) to S-GSAS/D614G (in blue) and the furin-cleaved S-RRAR/D614G ectodomain (in green) measured by ELISA. The assay format was the same as in Figure 2D. **C.** Overlay of the individual protomers in the 1-RBD-up structure and a protomer in the C3 symmetric 3-down-RBD structure shown in panel **A.** RBD-up chain with the S1 subunit colored by domain and the S2 subunit colored grey. RBD is colored red, NTD colored green, SD1 dark blue, SD2 orange and the linker between the NTD and RBD colored cyan. The down RBDs are colored salmon, the SD1 domains from the down RBD chains are colored light blue. The linker between the NTD and RBD in the down RBD chains are colored deep teal. Insets show zoomed-in views of individual domains similar to the depiction in Figure 4D. **D.** (left) The protomers of the 1-RBD-up structure of the furin-cleaved S-RRAR/D614G ectodomain superimposed using residues 908–1035 and colored by the color of their NTD as depicted in panel **A.** Zoomed-in views show region of the SD2 domain proximal to the NTD.

We compared the different protomers in the two structures by overlaying three protomers in the asymmetric 1-RBD-up structure and one protomer from the symmetric 3-RBD-down structure using residues 908–1035 (comprising the CH and HR1 regions) for superposition (Figure 7C). Similar to observations made with the S-GSAS/D614G S ectodomain structure, the RBD up/down motion in the furin-cleaved G614 S ectodomain was associated with a movement in the SD1 domain and in the region of the RBD-to-NTD linker that joined the SD1 β sheet (Figure 7C, **S8B**). As observed for S-GSAS/D614G, the SD2 domain showed little conformational change and formed a stable motif anchoring the mobile NTD and RBD domains. These observations reinforce the divergent roles of the SD1 and SD2 domain in RBD motion.

We next examined the region of the SD2 domain proximal to the NTD, and asked whether we could detect any structural changes in this region and if yes, could these be related to NTD motion. In the symmetric 3-RBD-down S ectodomain, all NTDs are identical, each stacking against the down RBD of the adjacent promoter. In the asymmetric 1-RBD-up structure, each NTD were distinct. To distinguish between these, we named the NTDs as follows: the NTD that was part of the protomer with the RBD in the up conformation was named NTD1. NTD1 stacked against a down RBD that contacted the up-RBD at one end and the second down-RBD at the other. The NTD2 stacked against a down-RBD that contacted a down-RBD at one end, and the NTD3 had the least amount of RBD contact by virtue of contacting the up-RBD (Figure 7A). Observing the NTD-proximal region on the SD2 domain (marked by a dotted square on Figure 7C) that also contacted the RBD-to-NTD linker, we noted shifts in the T602-606 loop between the different protomers. While the shifts were modest (with a maximal displacement of 2.2 Å), interestingly, identical trends were observed in the 1-RBD-up structures of the S-GSAS, S-GSAS/D614G and furin-cleaved S-GSAS/D614G S ectodomains, suggesting that this region of the SD2 domain responds to NTD motion and adopts a different conformation depending on the NTD environment (Figure 7D).

Thus, these data provide further evidence for allostery in the S protein, with changes in the SD2 domain impacting the RBD conformational dynamics. While the SD2 domain remains almost structurally invariant, we observe small but reproducible changes in SD2 loops in response to RBD/NTD movement suggesting that small changes in the SD2 region may translate to large motions in the RBD/NTD region.

## Discussion

Stabilized ectodomain constructs have proven to be useful tools to understand conformational dynamics of CoV S proteins. In particular, these have enabled high-resolution structural determination and atomic level understanding of the S ectodomain. They also are key components in vaccine development pipelines. The structural similarities in the S proteins of diverse CoVs have often enabled quick translation of structural rules and ideas from one CoV S ectodomain to another. Indeed, after the onset of the recent and ongoing COVID-19 pandemic, the SARS-CoV-2 S ectodomain could be rapidly stabilized and structurally characterized by exploiting its similarities with other CoVs and following strategies that had proved successful previously (Henderson et al., 2020; Pallesen et al., 2017; Wrapp et al., 2020). Some of these stabilization strategies, such as introduction of proline residues in the fusion subunit to prevent transition from pre-to post-fusion, have been successful in stabilizing the pre-fusion conformation of diverse class I fusion proteins including RSV F (Krarup et al., 2015), HIV-1 Env (Sanders et al., 2002), Ebola and Marburg GP (Rutten et al., 2020), influenza HA (Qiao et al., 1998) and Lassa GPC (Hastie et al., 2017). While the underlying hypothesis for the stabilization of the S ectodomain was that introduction of 2 proline residues at the junction of the CH and HR1 helices would arrest conformational transition to the post-fusion form, we found that even without the PP mutations, the SARS-CoV-2 S ectodomain retained its pre-fusion form. Not only so, even following furin cleavage the S ectodomain retained its pre-fusion conformation. These differences between the observed behavior of the SARS-CoV-2 S relative to other CoVs suggests that even though they retain similar overall topology and structural folds, there are differences between these CoVs that profoundly affect their structural and biological properties. Studying and accounting for these will be essential not only to understand SARS-CoV-2 but also to appreciate the nature and origin of these differences between CoVs for anticipating, preparing for and rapidly combating future CoV pandemics.

Viral surface proteins that are involved in receptor binding mediated cellular entry typically consist of flexible and moving parts that exhibit large conformational changes. While this conformational flexibility is necessary for function, structural checkpoints are required to prevent premature activation and destabilization or unfolding of the protein structure. Conformationally-silent structural islands provide the necessary stabilizing anchors for adjacent regions undergoing large motions. In this study we have identified the SD2 domain in the SARS-CoV-2 S protein as such a conformational anchor that is spatially interspersed between the highly mobile NTD and RBD regions, while itself remaining relatively invariant in its conformation. This conformational invariability of the SD2 subdomain is reminiscent of the beta sandwich structure in the HIV-1 envelope glycoprotein that connects and anchors a mobile layered architecture of the gp120 inner domain (Pancera et al., 2010). The conformationally invariant SD2 also serves to contain the movements of the RBD and NTD to the S1 subunit, such that the S2 subunit was unchanged between the various RBD “up” and “down” protomers (Figure 4 **and Figure S8**). This suggests a role for the SD2 domain in preventing premature triggering due to the stochastic up/down RBD motions in the SARS-CoV-2 S protein, as well as the importance of downstream events such as ACE2 receptor engagement and TMPRSS2 protease cleavage (Bestle et al., 2020; Hoffmann et al., 2020b; Matsuyama et al., 2020) in orchestrating the full extent of pre-to post-fusion transformation. In this study, we also assigned a key role to the N2R linker that connects the NTD to the RBD within a protomer. Rather than just being a connector, this 28-residue linker is also a modulator of conformational changes that are critical for receptor engagement. The linker contributes a beta strand to each of the SD1 and SD2 subdomains thus connecting all the structural domains in the S1 subunit.

In addition to the much discussed D614G mutation, the SD2 subdomain also houses the multibasic furin cleavage site that demarcates the S1 and S2 subunits. Furin cleavage is an essential processing step for the S protein and is necessary for viral infection and transmission (Hoffmann et al., 2020a; Shang et al., 2020). We provide evidence in this study that the D614G mutation enhances susceptibility of the SARS-CoV-2 S ectodomain to furin cleavage, thus raising the possibility that this is a contributor to increased fitness and transmissibility of D614G isolates.

### Limitations of this study

In this paper, we study the effect of the D614G mutation on RBD dynamics and susceptibility to furin cleavage. We find that the D614G mutation results in increased furin cleavage susceptibility, which could be responsible for the increased transmissibility of the SARS-CoV-2 with the D614G mutation. It is important to consider though that these results are obtained in the context of an engineered soluble construct and further studies are needed to understand if these effects translate to the native virion context.

## Acknowledgements

Cryo-EM data were collected at the National Center for Cryo-EM Access and Training (NCCAT) and the Simons Electron Microscopy Center located at the New York Structural Biology Center, supported by the NIH Common Fund Transformative High Resolution CryoElectron Microscopy program (U24 GM129539) and by grants from the Simons Foundation (SF349247) and NY State. This work was performed in part at the Duke University Shared Materials Instrumentation Facility (SMIF), a member of the North Carolina Research Triangle Nanotechnology Network (RTNN), which is supported by the National Science Foundation (award number ECCS-2025064) as part of the National Nanotechnology Coordinated Infrastructure (NNCI). We thank Ed Eng, Carolina Hernandez, Daija Bobe, Mark Walters and Holly Leddy for microscope alignments and assistance with cryo-EM data collection. This work was supported by an administrative supplement to NIH R01 AI145687 for coronavirus research (P.A. and R.C.H.) and NC State funding for COVID research (B.F.H). This study utilized the computational resources offered by Duke Research Computing (http://rc.duke.edu; NIH 1S10OD018164-01) at Duke University. We thank M. DeLong, C. Kneifel, M. Newton, V. Orlikowski, T. Milledge, and D. Lane from the Duke Office of Information Technology and Research Computing for assisting with setting up and maintaining the computing environment.

## Author contributions

S.M-C.G. and P.A. designed the study and wrote the manuscript with help from all authors. S.M-C.G. designed SARS-CoV-2 ectodomain constructs, expressed and purified proteins, performed biophysical and biochemical studies, performed NSEM experiments, determined and analyzed cryo-EM structures. K.J. expressed and purified proteins and performed protease cleavage experiments. S.M. and R.C.H. performed structural analysis related to the difference distance matrices. K.Mansouri collected and analyzed NSEM data. R.P. performed ELISA assays, K. Manne assisted with antigenicity and stability measurements. V.S. expressed and purified proteins. M.F.K. assisted with cryo-EM sample optimization. R.J.E. supervised NSEM studies. B.F.H supervised antigenicity measurements. P.A. supervised and led study and reviewed all data.

## STAR METHODS

### RESOURCE AVAILABILITY

#### Lead Contact and Materials Availability

Further information and requests for resources and reagents should be directed to Priyamvada Acharya.

#### Data and Code Availability

Cryo-EM reconstructions and atomic models generated during this study are available at wwPDB and EMBD (https://www.rcsb.org; http://emsearch.rutgers.edu) under the accession codes PDB IDs 7KDG, 7KDH, 7KDK, 7KDL, 7KDI, 7KDJ, 7KE4, 7KE6, 7KE7, 7KE8, 7KE9, 7KEA, 7KEB, 7KEC and EMDB IDs EMDB-22821, EMD-22822, EMD-22825, EMD-22826, EMD-22823, EMD-22824, EMD-22831, EMD-22832, EMD-22833, EMD-22834, EMD-22835, EMD-22835, EMD-22837, EMD-22838.

### EXPERIMENTAL MODEL AND SUBJECT DETAILS

#### Cell culture

Gibco FreeStyle 293-F cells (embryonal, human kidney) were incubated at 37°C and 9% CO_2_ in a humidified atmosphere. Cells were incubated in FreeStyle 293 Expression Medium (Gibco) with agitation at 120 rpm. Plasmids were transiently transfected into cells using Turbo293 (SpeedBiosystems) and incubated at 37 °C, 9% CO2, 120 rpm for 6 days. On the day following transfection, HyClone CDM4HEK293 media (Cytiva, MA) was added to the cells.

Antibodies were produced in Expi293 cells (embryonal, human kidney). Cells were incubated in Expi293 Expression Medium at 37°C, 120 rpm and 8% CO_2_ in a humidified atmosphere. Plasmids were transiently transfected into cells using the ExpiFectamine 293 Transfection Kit and protocol (Gibco).

### METHOD DETAILS

#### Plasmids

All genes in this study were synthesized and sequenced by GeneImmune Biotechnology (Rockville, MD). The SARS-CoV-2 spike protein ectodomain constructs used comprised the protein residues 1–1208 (GenBank: MN908947) with or without the D614G mutation, with or without the furin cleavage site RRAR (residue 682–685) mutated to GSAS or LEVLFQGP (HRV3C protease site), a C-terminal T4 fibritin trimerization motif, a C-terminal HRV3C protease cleavage site (except for the constructs where the furin site was mutated to an HRV3C site), a TwinStrepTag and an 8XHisTag. All spike ectodomain constructs were cloned into the mammalian expression vector pαH (Wrapp et al., 2020). For the ACE-2 construct, the C-terminus was fused a human Fc region.

#### Protein purification

Spike ectodomains were harvested from filtered and concentrated supernatant using StrepTactin resin (IBA) and further purified by SEC using a Superose 6 10/300 GL Increase column preequilibrated in 2mM Tris, pH 8.0, 200 mM NaCl, 0.02% sodium azide. All protein purification steps were performed at room temperature in a single day. The purified proteins were flash frozen and stored at −80 °C in single-use aliquots. Each aliquot were thawed by incubation (∼20 min) at 37 °C before use.

Antibodies were produced in Expi293F cells and purified by Protein A affinity. ACE-2 with human Fc tag was purified by Protein A affinity chromatography.

#### Negative-stain electron microscopy

Samples were diluted to 100 μg/ml in 20 mM HEPES pH 7.4, 150 mM NaCl, 5% glycerol, 7.5 mM glutaraldehyde and incubated for 5 minutes before quenching the glutaraldehyde by the addition of 1 M Tris (to a final concentration of 75 mM) and 5 minutes incubation. A 5-μl drop of sample was then applied to a glow-discharged carbon-coated grid for 10–15 seconds, blotted, stained with 2% uranyl formate, blotted and air-dried. Images were obtained using a Philips EM420 electron microscope at 120 kV, 82,000× magnification, and a 4.02 Å pixel size. The RELION (Scheres, 2012) program was used for particle picking, 2D and 3D class averaging.

#### Differential scanning fluorimetry

DSF assay was performed using Tycho NT. 6 (NanoTemper Technologies). Spike ectodomains were diluted to approximatively 0.15 mg/ml. Intrinsic fluorescence was measured at 330 nm and 350 nm while the sample was heated from 35 to 95 °C at a rate of 30°C/min. The ratio of fluorescence (350/330 nm) and inflection temperatures (Ti) were calculated by the Tycho NT. 6 apparatus.

#### ELISA assays

Spike samples were pre-incubated at different temperatures then tested for antibody- or ACE-2-binding in ELISA assays as previously described (Edwards et al., 2020). Assays were run in two formats. In the first format antibodies or ACE2 protein were coated on 384-well plates at 2 μg/ml overnight at 4°C, washed, blocked and followed by two-fold serially diluted spike protein starting at 25 μg/mL. Binding was detected with polyclonal anti-SARS-CoV-2 spike rabbit serum (developed in our lab), followed by goat anti-rabbit-HRP and TMB substrate. Absorbance was read at 450 nm. In the second format, serially diluted spike protein was bound in individual wells of 384-well plates, which were previously coated with streptavidin at 2 μg/mL and blocked. Proteins were incubated at room temperature for 1 hour, washed, then human mAbs were added at 10 μg/ml. Antibodies were incubated at room temperature for 1 hour, washed and binding detected with goat anti-human-HRP and TMB substrate.

#### Cryo-EM

Purified SARS-CoV-2 spike preparations were diluted to a concentration of ∼1.5 mg/mL in 2 mM Tris pH 8.0, 200 mM NaCl and 0.02% NaN_3_. A 2.5-μL drop of protein was deposited on a Quantifoil-1.2/1.3 grid that had been glow discharged for 10 seconds in a PELCO easiGlow™ Glow Discharge Cleaning System. After a 30 seconds incubation in >95% humidity, excess protein was blotted away for 2.5 seconds before being plunge frozen into liquid ethane using a Leica EM GP2 plunge freezer (Leica Microsystems). Frozen grids were imaged in a Titan Krios (Thermo Fisher) equipped with a K3 detector (Gatan).

### QUANTIFICATION AND STATISTICAL ANALYSIS

No statistical analysis were performed in this study.

**Key Resources Table**

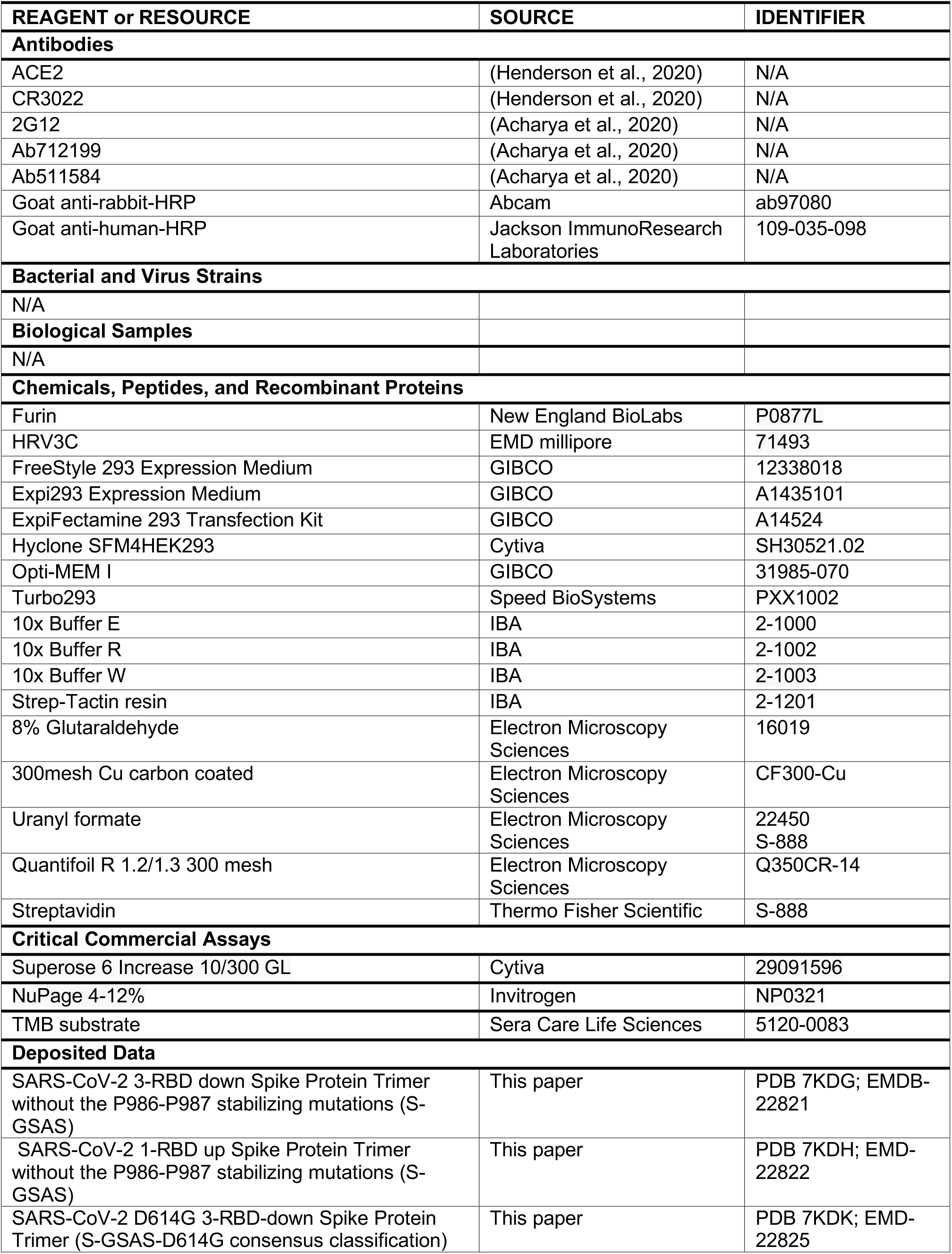
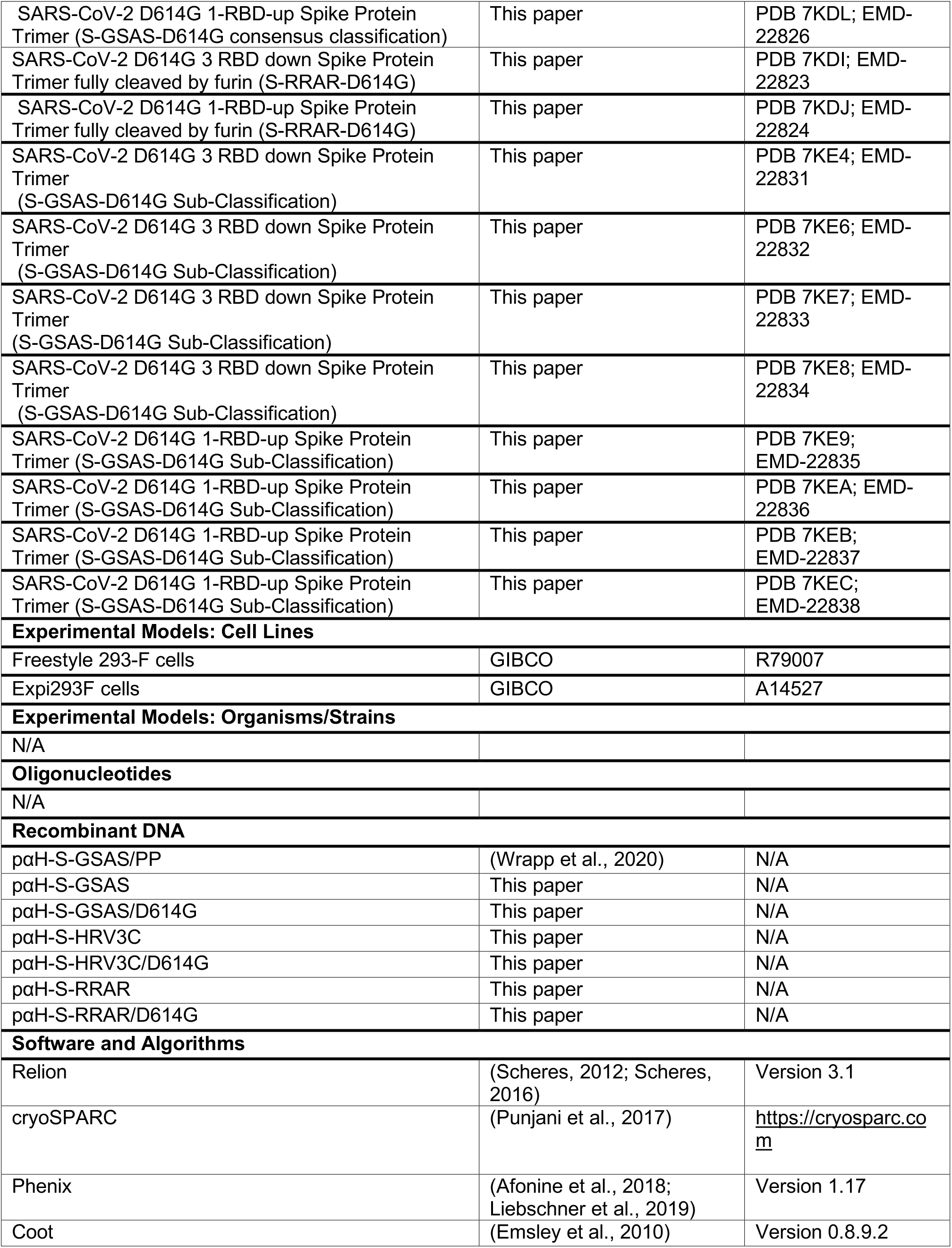
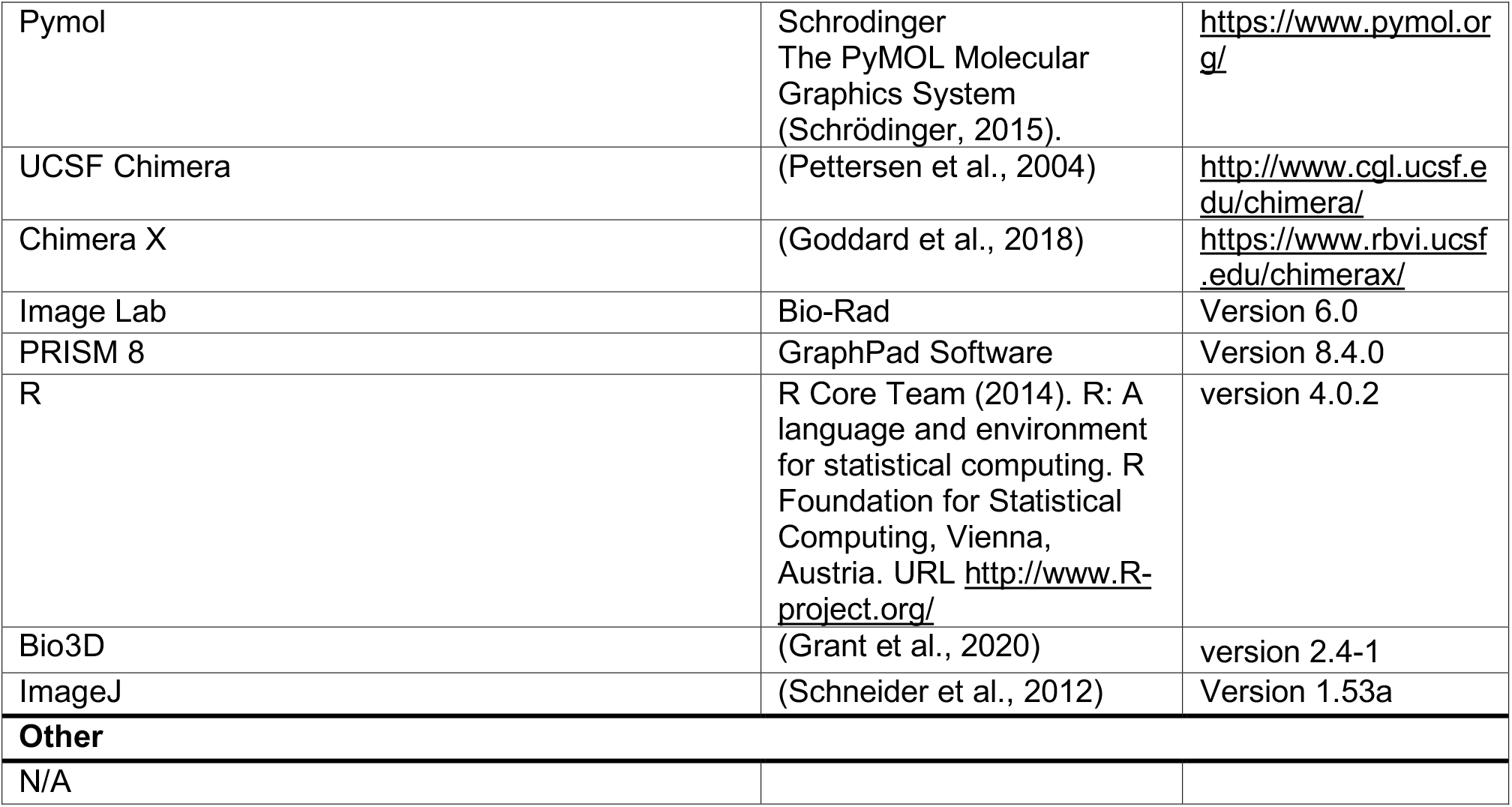

| REAGENT or RESOURCE | SOURCE | IDENTIFIER |
| --- | --- | --- |
| **Antibodies** |  |  |
| ACE2 | (Henderson et al., 2020) | N/A |
| CR3022 | (Henderson et al., 2020) | N/A |
| 2G12 | (Acharya et al., 2020) | N/A |
| Ab712199 | (Acharya et al., 2020) | N/A |
| Ab511584 | (Acharya et al., 2020) | N/A |
| Goat anti-rabbit-HRP | Abcam | ab97080 |
| Goat anti-human-HRP | Jackson ImmunoResearch Laboratories | 109-035-098 |
| **Bacterial and Virus Strains** |  |  |
| N/A |  |  |
| **Biological Samples** |  |  |
| N/A |  |  |
| **Chemicals, Peptides, and Recombinant Proteins** |  |  |
| Furin | New England BioLabs | P0877L |
| HRV3C | EMD millipore | 71493 |
| FreeStyle 293 Expression Medium | GIBCO | 12338018 |
| Expi293 Expression Medium | GIBCO | A1435101 |
| ExpiFectamine 293 Transfection Kit | GIBCO | A14524 |
| Hyclone SFM4HEK293 | Cytiva | SH30521.02 |
| Opti-MEM I | GIBCO | 31985-070 |
| Turbo293 | Speed BioSystems | PXX1002 |
| 10x Buffer E | IBA | 2-1000 |
| 10x Buffer R | IBA | 2-1002 |
| 10x Buffer W | IBA | 2-1003 |
| Strep-Tactin resin | IBA | 2-1201 |
| 8% Glutaraldehyde | Electron Microscopy Sciences | 16019 |
| 300mesh Cu carbon coated | Electron Microscopy Sciences | CF300-Cu |
| Uranyl formate | Electron Microscopy Sciences | 22450<br>S-888 |
| Quantifoil R 1.2/1.3 300 mesh | Electron Microscopy Sciences | Q350CR-14 |
| Streptavidin | Thermo Fisher Scientific | S-888 |
| **Critical Commercial Assays** |  |  |
| Superose 6 Increase 10/300 GL | Cytiva | 29091596 |
| NuPage 4-12% | Invitrogen | NP0321 |
| TMB substrate | Sera Care Life Sciences | 5120-0083 |
| **Deposited Data** |  |  |
| SARS-CoV-2 3-RBD down Spike Protein Trimer without the P986-P987 stabilizing mutations (S-GSAS) | This paper | PDB 7KDG; EMD-22821 |
| SARS-CoV-2 1-RBD up Spike Protein Trimer without the P986-P987 stabilizing mutations (S-GSAS) | This paper | PDB 7KDH; EMD-22822 |
| SARS-CoV-2 D614G 3-RBD-down Spike Protein Trimer (S-GSAS-D614G consensus classification) | This paper | PDB 7KDK; EMD-22825 |
| SARS-CoV-2 D614G 1-RBD-up Spike Protein Trimer (S-GSAS-D614G consensus classification) | This paper | PDB 7KDL; EMD-22826 |
| SARS-CoV-2 D614G 3 RBD down Spike Protein Trimer fully cleaved by furin (S-RRAR-D614G) | This paper | PDB 7KDI; EMD-22823 |
| SARS-CoV-2 D614G 1-RBD-up Spike Protein Trimer fully cleaved by furin (S-RRAR-D614G) | This paper | PDB 7KDJ; EMD-22824 |
| SARS-CoV-2 D614G 3 RBD down Spike Protein Trimer (S-GSAS-D614G Sub-Classification) | This paper | PDB 7KE4; EMD-22831 |
| SARS-CoV-2 D614G 3 RBD down Spike Protein Trimer (S-GSAS-D614G Sub-Classification) | This paper | PDB 7KE6; EMD-22832 |
| SARS-CoV-2 D614G 3 RBD down Spike Protein Trimer (S-GSAS-D614G Sub-Classification) | This paper | PDB 7KE7; EMD-22833 |
| SARS-CoV-2 D614G 3 RBD down Spike Protein Trimer (S-GSAS-D614G Sub-Classification) | This paper | PDB 7KE8; EMD-22834 |
| SARS-CoV-2 D614G 1-RBD-up Spike Protein Trimer (S-GSAS-D614G Sub-Classification) | This paper | PDB 7KE9; EMD-22835 |
| SARS-CoV-2 D614G 1-RBD-up Spike Protein Trimer (S-GSAS-D614G Sub-Classification) | This paper | PDB 7KEA; EMD-22836 |
| SARS-CoV-2 D614G 1-RBD-up Spike Protein Trimer (S-GSAS-D614G Sub-Classification) | This paper | PDB 7KEB; EMD-22837 |
| SARS-CoV-2 D614G 1-RBD-up Spike Protein Trimer (S-GSAS-D614G Sub-Classification) | This paper | PDB 7KEC; EMD-22838 |
| **Experimental Models: Cell Lines** |  |  |
| Freestyle 293-F cells | GIBCO | R79007 |
| Expi293F cells | GIBCO | A14527 |
| **Experimental Models: Organisms/Strains** |  |  |
| N/A |  |  |
| **Oligonucleotides** |  |  |
| N/A |  |  |
| **Recombinant DNA** |  |  |
| paH-S-GSAS/PP | (Wrapp et al., 2020) | N/A |
| paH-S-GSAS | This paper | N/A |
| paH-S-GSAS/D614G | This paper | N/A |
| paH-S-HRV3C | This paper | N/A |
| paH-S-HRV3C/D614G | This paper | N/A |
| paH-S-RRAR | This paper | N/A |
| paH-S-RRAR/D614G | This paper | N/A |
| **Software and Algorithms** |  |  |
| Relion | (Scheres, 2012; Scheres, 2016) | Version 3.1 |
| cryoSPARC | (Punjani et al., 2017) | https://cryosparc.com |
| Phenix | (Afonine et al., 2018; Liebschner et al., 2019) | Version 1.17 |
| Coot | (Emsley et al., 2010) | Version 0.8.9.2 |
| Pymol | Schrodinger<br>The PyMOL Molecular Graphics System<br>(Schrödinger, 2015). | https://www.pymol.org/ |
| UCSF Chimera | (Pettersen et al., 2004) | http://www.cgl.ucsf.edu/chimera/ |
| Chimera X | (Goddard et al., 2018) | https://www.rbvi.ucsf.edu/chimerax/ |
| Image Lab | Bio-Rad | Version 6.0 |
| PRISM 8 | GraphPad Software | Version 8.4.0 |
| R | R Core Team (2014). R: A language and environment for statistical computing. R Foundation for Statistical Computing, Vienna, Austria. URL http://www.R-project.org/ | version 4.0.2 |
| Bio3D | (Grant et al., 2020) | version 2.4-1 |
| ImageJ | (Schneider et al., 2012) | Version 1.53a |
| **Other** |  |  |
| N/A |  |  |

