## Supplemental Data for "D614G mutation alters SARS-CoV-2 spike conformational dynamics and protease cleavage susceptibility at the S1/S2 junction"

**Table S1. Cryo-EM data collection and refinements statistics for the S-GSAS, S-GSAS/D614G and furin cleaved S-RRAR/D614G SARS-CoV-2 spike ectodomains.**

| | S-GSAS | | S-GSAS(D614G) | | S-RRAR(D614G) | |
|---|---|---|---|---|---|---|
| | 3-down | 1-up | 3-down consensus | 1-up consensus | 3-down | 1-up |
| PDB code | 7KDG | 7KDH | 7KDK | 7KDL | 7KDI | 7KDJ |
| Data collection and processing | | | | | | |
| Microscope | FEI Titan Krios | | | | | |
| Detector | Gatan K3 | | | | | |
| Magnification | 81,000 | | 81,000 | | 81,000 | |
| Voltage (kV) | 300 | | 300 | | 300 | |
| Electron exposure (e–/Å2) | 65.24 | | 51.8 | | 65.94 | |
| Defocus range (μm) | 0.8-2.5 | | 0.40-2.94 | | 0.38-2.88 | |
| Pixel size (Å) | 1.069 | | 1.058 | | 1.058 | |
| Reconstruction software | cryoSparc | | | | | |
| Symmetry imposed | C3 | C1 | C3 | C1 | C1 | C1 |
| Initial particle images (no.) | 2,566,724 | | 2,343,150 | | 479,208 | |
| Final particle images (no.) | 581,495 | 175,529 | 782,485 | 613,271 | 224,310 | 99,165 |
| Map resolution (Å) | 3.01 | 3.33 | 2.8 | 2.96 | 3.26 | 3.49 |
| FSC threshold | 0.143 | 0.143 | 0.143 | 0.143 | 0.143 | 0.143 |
| Refinement | | | | | | |
| Initial model used | 6VXX | 6VYB | 6VXX | 6VYB | 6VXX | 6VYB |
| Model resolution (Å) | 3.01 | 3.33 | 2.8 | 2.96 | 3.26 | 3.49 |
| FSC threshold | 0.143 | 0.143 | 0.143 | 0.143 | 0.143 | 0.143 |
| Map sharpening B factor ($Å^2$) | -129.3 | -101.3 | -121.5 | -106.9 | -104.2 | -94.8 |
| Model composition | | | | | | |
| Nonhydrogen atoms | 23,700 | 22,698 | 23,688 | 22,365 | 23,688 | 22,365 |
| Protein residues | 2,916 | 2,891 | 2,916 | 2,875 | 2,916 | 2,875 |
| R.m.s. deviations | | | | | | |
| Bond lengths (Å) | 0.013 | 0.012 | 0.013 | 0.012 | 0.012 | 0.012 |
| Bond angles (°) | 1.845 | 1.838 | 1.87 | 1.835 | 1.789 | 1.757 |
| Validation | | | | | | |
| MolProbity score | 1.03 | 1.13 | 0.98 | 0.96 | 0.97 | 1.03 |
| Clashscore | 0.28 | 0.43 | 0.3 | 0.09 | 0.15 | 0.23 |
| Poor rotamers (%) | 0.31 | 0.56 | 0.12 | 0.27 | 0.27 | 0.44 |
| Ramachandran plot | | | | | | |
| Favored (%) | 93.46 | 92.3 | 94.66 | 93.6 | 93.81 | 93.21 |
| Allowed (%) | 6.22 | 7.31 | 5.24 | 6.11 | 6.12 | 6.4 |
| Disallowed (%) | 0.32 | 0.39 | 0.11 | 0.29 | 0.07 | 0.39 |

**Table S1 continued. Cryo-EM data collection and refinements statistics for the S-GSAS, S-GSAS/D614G and S-RRAR/D614G SARS-CoV-2 spike protein ectodomains.**

| | S-GSAS(D614G) | | | | | | | |
|---|---|---|---|---|---|---|---|---|
| | 3-down | | | | 1-up | | | |
| PDB code | 7KE4 | 7KE6 | 7KE7 | 7KE8 | 7KE9 | 7KEA | 7KEB | 7KEC |
| Data collection and processing | | | | | | | | |
| Magnification | 81,000 | | | | | | | |
| Voltage (kV) | 300 | | | | | | | |
| Electron exposure (e–/Å$^2$) | 51.8 | | | | | | | |
| Defocus range (μm) | 0.40-2.94 | | | | | | | |
| Pixel size (Å) | 1.058 | | | | | | | |
| Symmetry imposed | C1 | | | | | | | |
| Initial particle images (no.) | 2,343,150 | | | | | | | |
| Final particle images (no.) | 182,326 | 196,173 | 157,254 | 133,373 | 287,765 | 145,566 | 127,429 | 58,580 |
| Map resolution (Å) | 3.21 | 3.1 | 3.32 | 3.26 | 3.08 | 3.33 | 3.48 | 3.84 |
| FSC threshold | 0.143 | 0.143 | 0.143 | 0.143 | 0.143 | 0.143 | 0.143 | 0.143 |
| Refinement | | | | | | | | |
| Initial model used | 6VXX | 6VXX | 6VXX | 6VXX | 6VYB | 6VYB | 6VYB | 6VYB |
| Model resolution (Å) | 3.21 | 3.1 | 3.32 | 3.26 | 3.08 | 3.33 | 3.48 | 3.84 |
| FSC threshold | 0.143 | 0.143 | 0.143 | 0.143 | 0.143 | 0.143 | 0.143 | 0.143 |
| Map sharpening B factor (Å$^2$) | -95.3 | -98.8 | -92.8 | -91.2 | -104.5 | -88.9 | -86.2 | -81 |
| Model composition | | | | | | | | |
| Nonhydrogen atoms | 23,684 | 23,688 | 23,688 | 23,684 | 22,365 | 22,365 | 22,365 | 22,337 |
| Protein residues | 2,916 | 2,916 | 2,916 | 2,916 | 2,875 | 2,875 | 2,875 | 2,875 |
| R.m.s. deviations | | | | | | | | |
| Bond lengths (Å) | 0.013 | 0.013 | 0.013 | 0.013 | 0.012 | 0.012 | 0.012 | 0.012 |
| Bond angles (°) | 1.845 | 1.838 | 1.81 | 1.836 | 1.813 | 1.893 | 1.856 | 1.845 |
| Validation | | | | | | | | |
| MolProbity score | 1.02 | 1 | 1.1 | 1.13 | 0.95 | 1.23 | 1.3 | 1.58 |
| Clashscore | 0.13 | 0.15 | 0.34 | 0.41 | 0.11 | 0.64 | 0.83 | 0.74 |
| Poor rotamers (%) | 1.06 | 1.02 | 0.08 | 1.1 | 0.04 | 1.24 | 0.13 | 2.44 |
| Ramachandran plot | | | | | | | | |
| Favored (%) | 92.93 | 93.25 | 92.23 | 92.9 | 93.99 | 92.42 | 90.31 | 89.92 |
| Allowed (%) | 6.72 | 6.47 | 7.49 | 6.82 | 5.79 | 7.15 | 9.08 | 9.05 |
| Disallowed (%) | 0.35 | 0.28 | 0.28 | 0.28 | 0.21 | 0.43 | 0.61 | 1.04 |

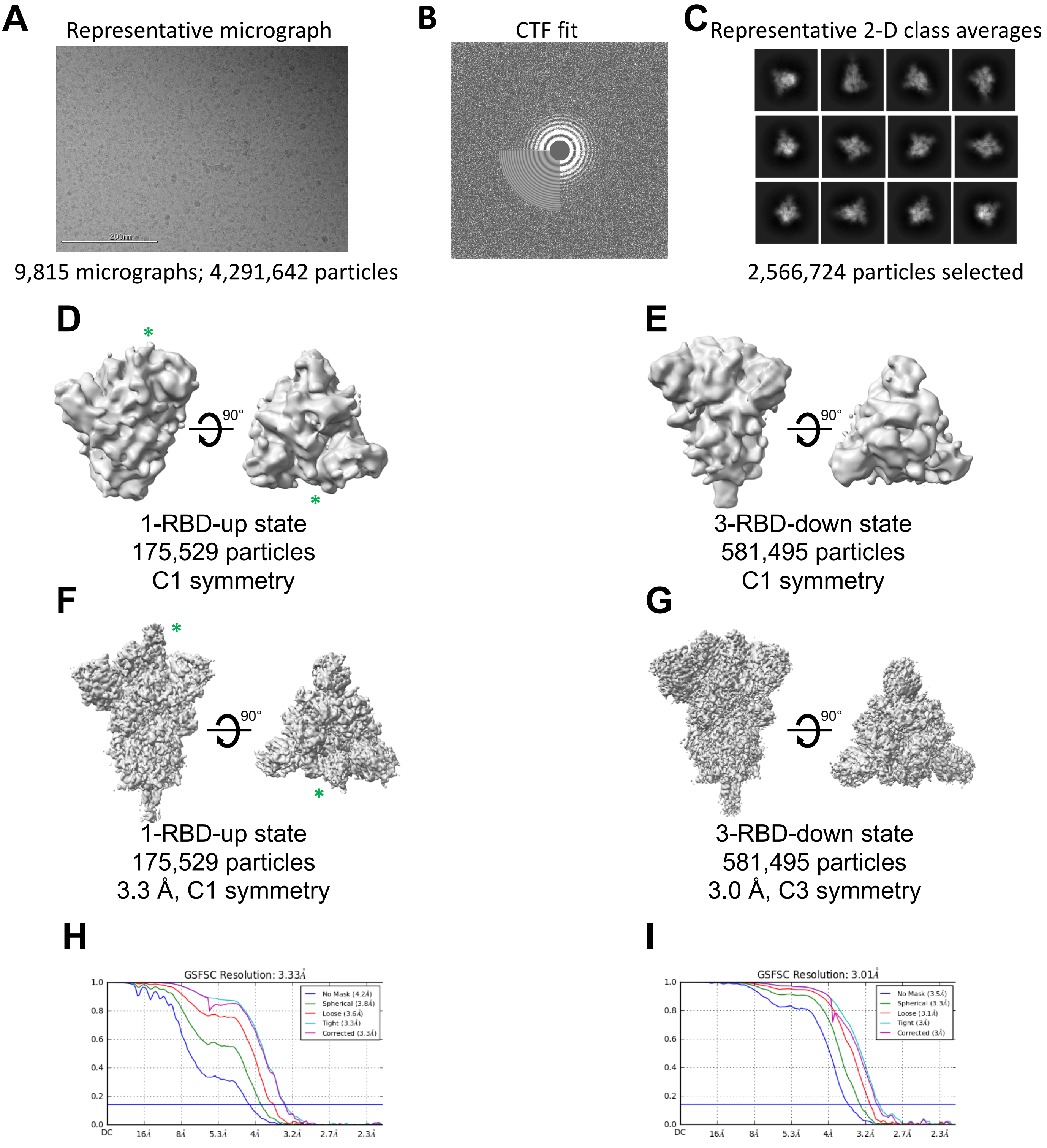


**Figure S1. Cryo-EM data processing details for the S-GSAS ectodomain. A.** Representative micrograph. **B.** CTF fit. **C.** Representative 2D class averages. **D-E.** *Ab initio* reconstructions for the **D.** 1-RBD-up spike (the RBD in the up position is identified by an asterisk) and **E.** 3-RBD-down spike. **F-G.** Refined maps for the **F.** 1-RBD-up spike (the RBD in the up position is identified by an asterisk) and **G** 3-RBD-down spike. **H-I.** Fourier shell correlation curves for the **H.** 1-RBD-up spike reconstruction and **I.** 3-RBD-down spike reconstruction.

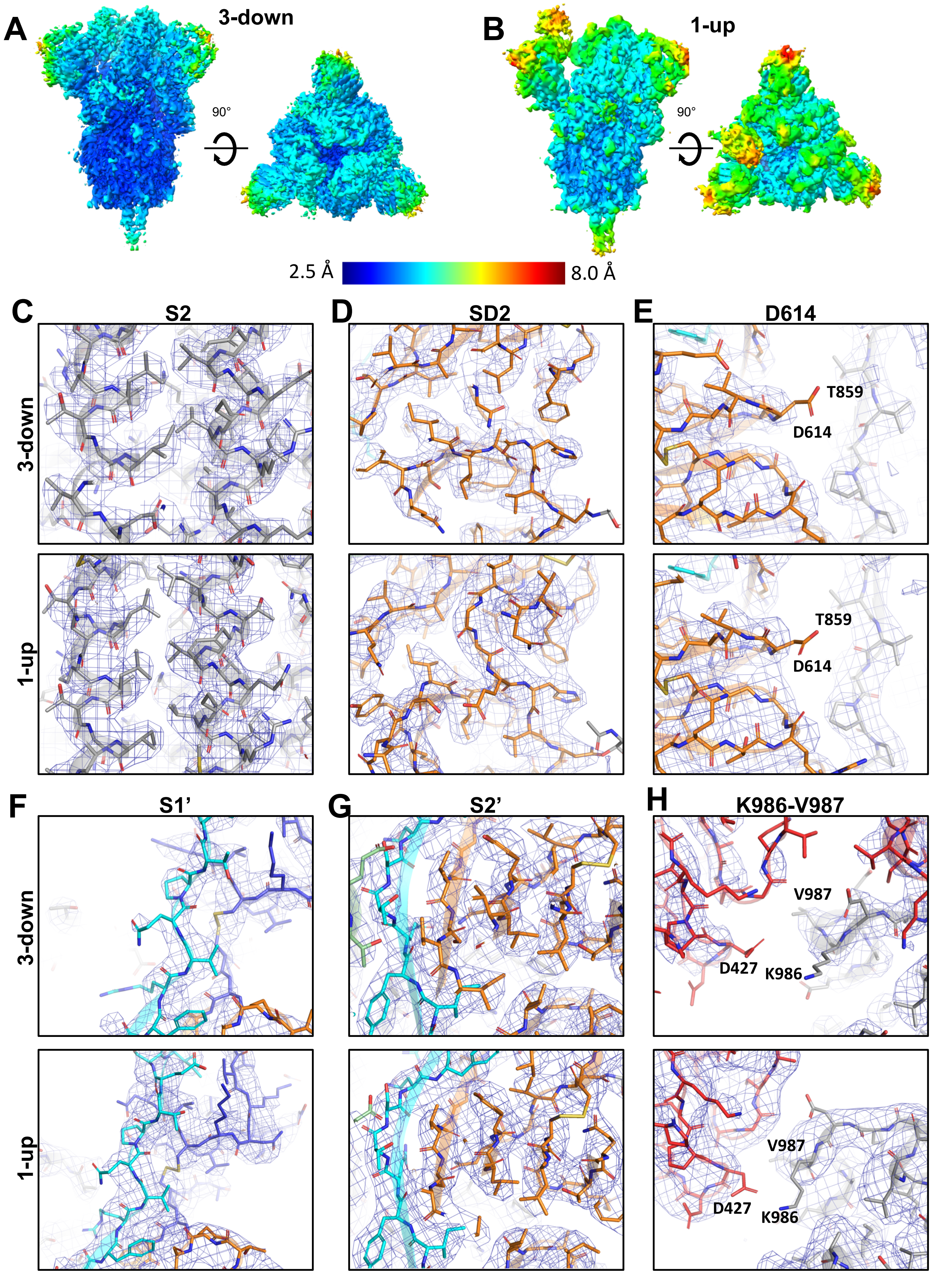

A
3-down
90°
B
1-up
90°
2.5 Å
8.0 Å
C
S2
D
SD2
E
D614
T859
D614
T859
D614
3-down
1-up
F
S1'
G
S2'
H
K986-V987
V987
D427
K986
V987
D427
K986
3-down
1-up

**Figure S2. Local cryo-EM map resolutions for the S-GSAS ectodomain. A-B.** Refined cryo-EM maps colored by local resolution for the **A.** 3-RBD-down and **B.** 1-RDB-up spikes. **C-H.** Zoomed-in images showing representative views of the **C.** S2 subunit, **D.** SD2 subdomain, **E.** region around the D614 mutation, **F.** S1' “super” subdomain **G.** S2' “super” subdomain and **H.** K986-V987 regions in the 3-RBD-down (top) and 1-RDB-up spike structures (bottom). The cryo-EM map is shown as a blue mesh and the fitted model is in cartoon representation, with residues shown as sticks.

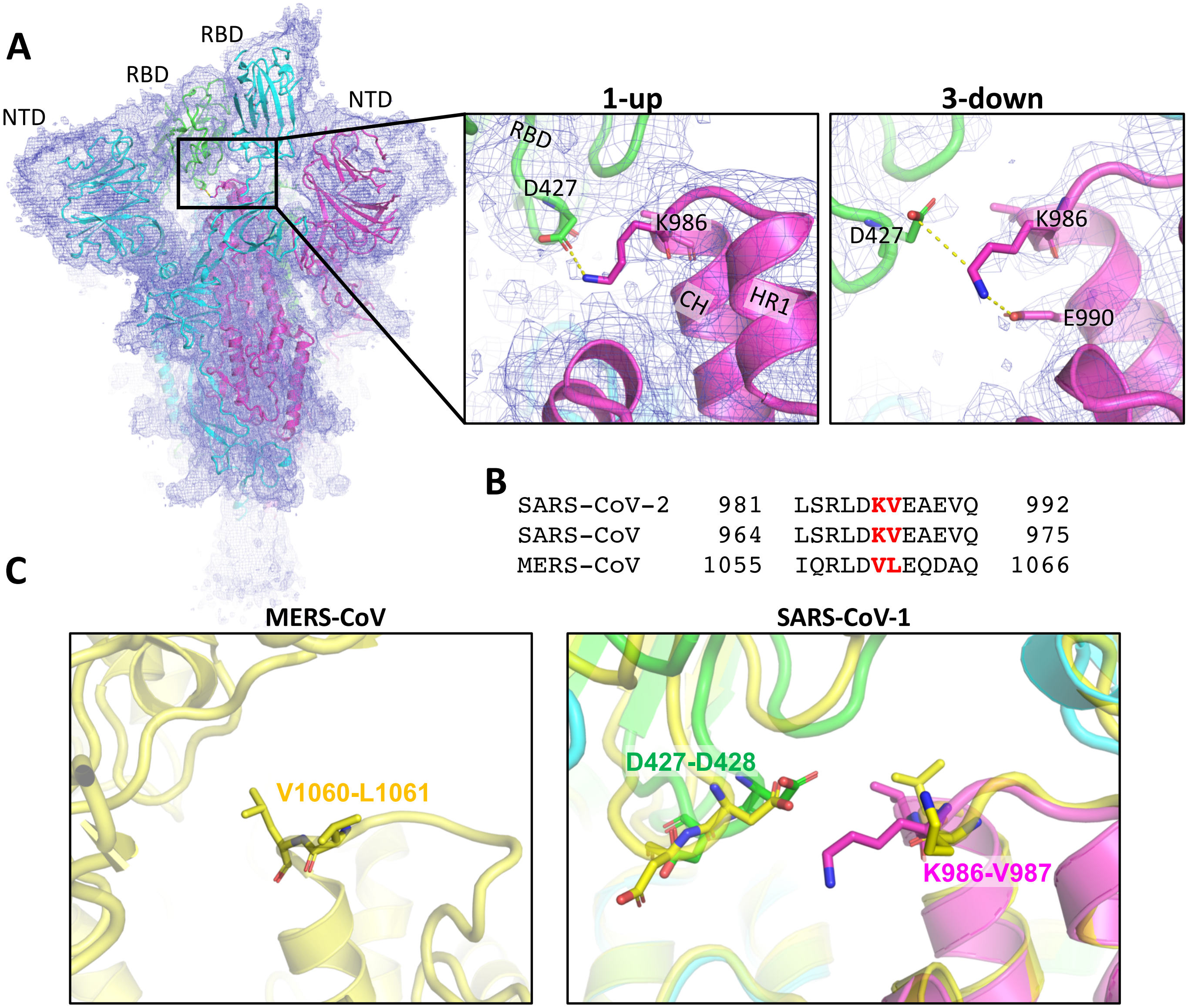


**Figure S3. Structural comparison of residues 986-987 in the SARS-CoV-2 S ectodomain with the MERS and SARS-CoV-1 S ectodomains. A.** Cryo-EM reconstruction map of the 1-RBD-up spike of the SARS-CoV-2 S-GSAS ectodomain colored by chains, with insets showing zoomed-in view of residues K986-V987 in the 1-RBD-up and 3-RBD-down spike structures. The cryo-EM map is shown as a blue mesh and the fitted model in cartoon representation, with key residues shown as sticks. **B.** Sequence alignment of residues 981-992 of SARS-CoV-2 and corresponding residues of SARS-CoV-1 and MERS spike proteins. **C.** Magnified view of one protomer showing residues V1060 and L1061 from MERS (PDB 5X5C; in yellow) and SARS-CoV-1 (PDB 5X58; in yellow) overlayed with SARS-CoV-2 S-GSAS ectodomain (colored as in **A**).

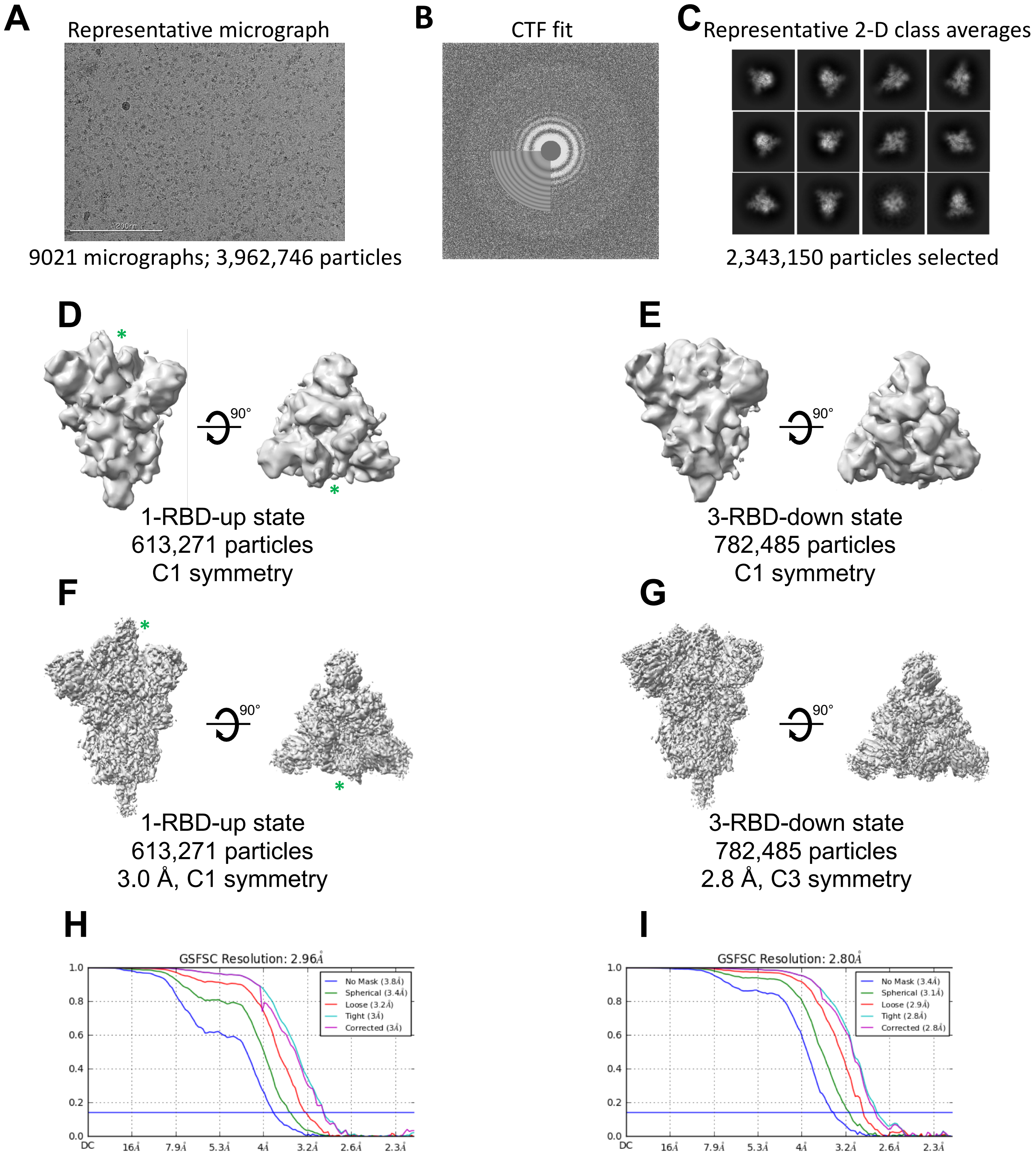


**Figure S4. Cryo-EM data processing details for S-GSAS/D614G consensus structures in Figure 3C. A.** Representative micrograph. **B.** CTF fit. **C.** Representative 2D class averages. **D-E.** *Ab initio* reconstructions for the **D.** consensus 1-RBD-up state (the RBD in the up position is identified by an asterisk) and **E.** consensus 3-RBD-down state. **F-G.** Refined maps for the **F.** consensus 1-RBD-up state (the RBD in the up position is identified by an asterisk) and **G.** consensus 3-RBD-down state. **H-I.** Fourier shell correlation curves for the **H.** consensus 1-RBD-up state and **I.** consensus 3-RBD-down state.

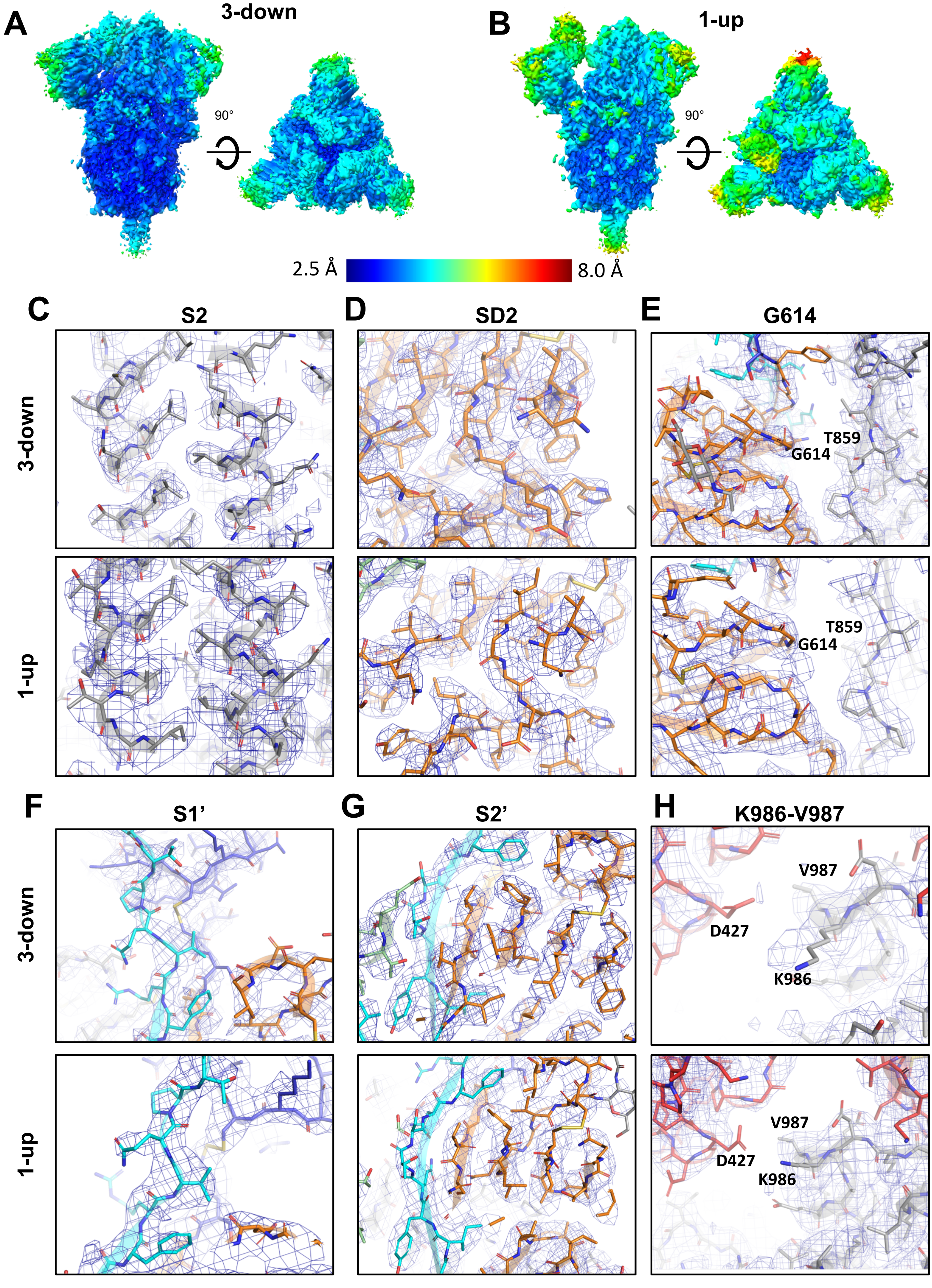
A
3-down
90°
B
1-up
90°
2.5 Å
8.0 Å
C
S2
D
SD2
E
G614
3-down
T859
G614
1-up
T859
G614
F
S1'
G
S2'
H
K986-V987
3-down
V987
D427
K986
1-up
V987
D427
K986

**Figure S5. Local map resolutions for S-GSAS/D614G ectodomain. A-B.** Refined cryo-EM maps colored by local resolution for the **A.** consensus 3-RBD-down and **B.** consensus 1-RDB-up states. **C-H.** Zoomed-in images showing representative views of the **C.** S2 subunit, **D.** SD2 subdomain, **E.** region around the D614 mutation, **F.** S1' "super" subdomain **G.** S2' "super" subdomain and **H.** K986-V987 regions in the 3-RBD-down (top) and 1-RDB-up spike structures (bottom). The cryo-EM map is shown as a blue mesh and the fitted model is in cartoon representation, with residues shown as sticks.

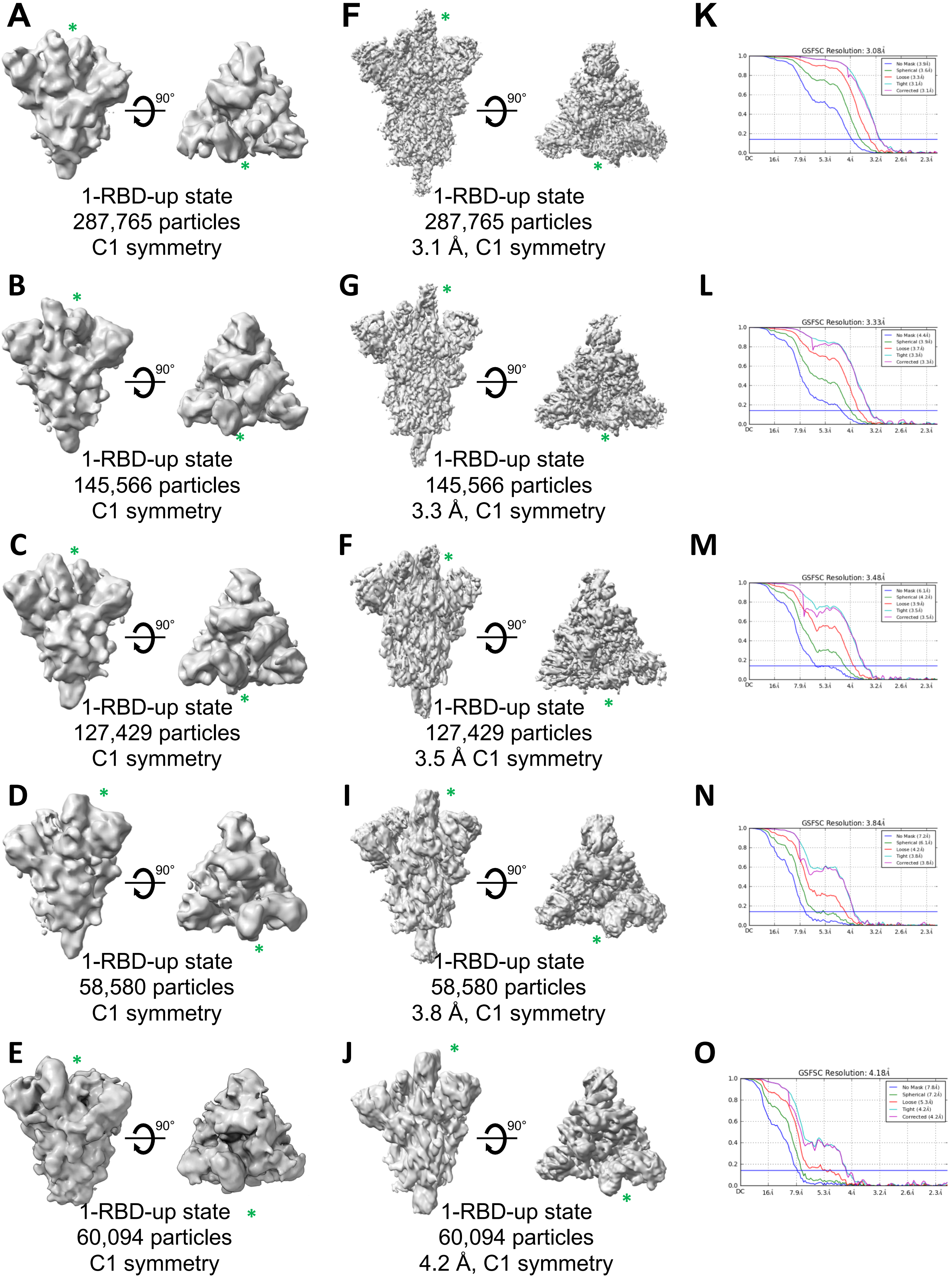
A
90°
1-RBD-up state
287,765 particles
C1 symmetry
F
90°
1-RBD-up state
287,765 particles
3.1 Å, C1 symmetry
K
GSFSC Resolution: 3.08Å
B
90°
1-RBD-up state
145,566 particles
C1 symmetry
G
90°
1-RBD-up state
145,566 particles
3.3 Å, C1 symmetry
L
GSFSC Resolution: 3.33Å
C
90°
1-RBD-up state
127,429 particles
C1 symmetry
F
90°
1-RBD-up state
127,429 particles
3.5 Å C1 symmetry
M
GSFSC Resolution: 3.48Å
D
90°
1-RBD-up state
58,580 particles
C1 symmetry
I
90°
1-RBD-up state
58,580 particles
3.8 Å, C1 symmetry
N
GSFSC Resolution: 3.84Å
E
90°
1-RBD-up state
60,094 particles
C1 symmetry
J
90°
1-RBD-up state
60,094 particles
4.2 Å, C1 symmetry
O
GSFSC Resolution: 4.18Å

**Figure S6. Cryo-EM data processing details for S-GSAS/D614G 1-up subpopulations from Figure 3D. A-E.** *Ab initio* reconstructions for the 1-RBD-up subpopulations (the RBD in the up position is identified by an asterisk). **F-J.** Refined maps for the 1-RBD-up subpopulations (the RBD in the up position is identified by an asterisk). **K-O.** Fourier shell correlation curves for the 1-RBD-up subpopulations.

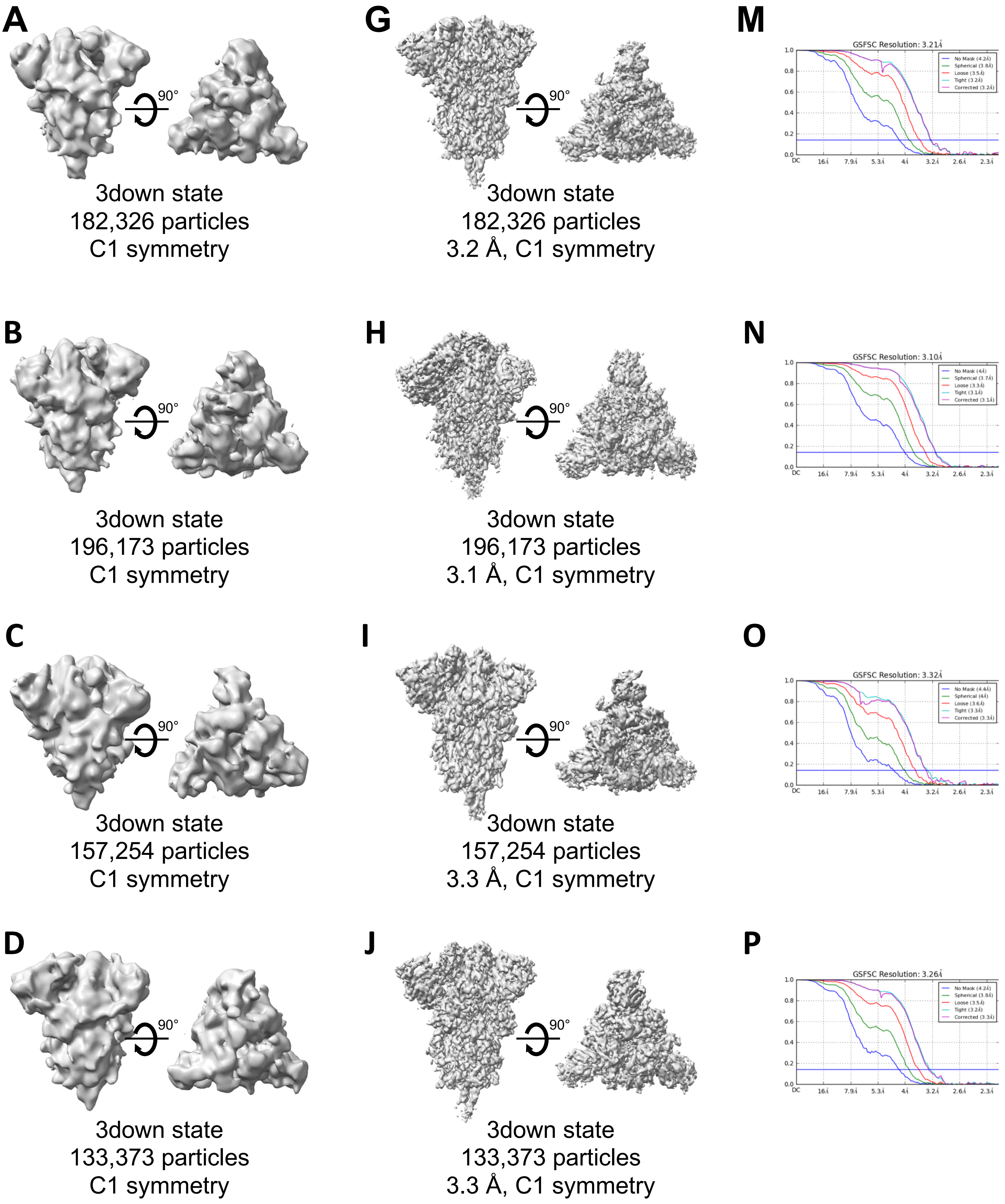
A
90°
3down state
182,326 particles
C1 symmetry
G
90°
3down state
182,326 particles
3.2 Å, C1 symmetry
M
GSFSC Resolution: 3.21Å
B
90°
3down state
196,173 particles
C1 symmetry
H
90°
3down state
196,173 particles
3.1 Å, C1 symmetry
N
GSFSC Resolution: 3.10Å
C
90°
3down state
157,254 particles
C1 symmetry
I
90°
3down state
157,254 particles
3.3 Å, C1 symmetry
O
GSFSC Resolution: 3.32Å
D
90°
3down state
133,373 particles
C1 symmetry
J
90°
3down state
133,373 particles
3.3 Å, C1 symmetry
P
GSFSC Resolution: 3.26Å

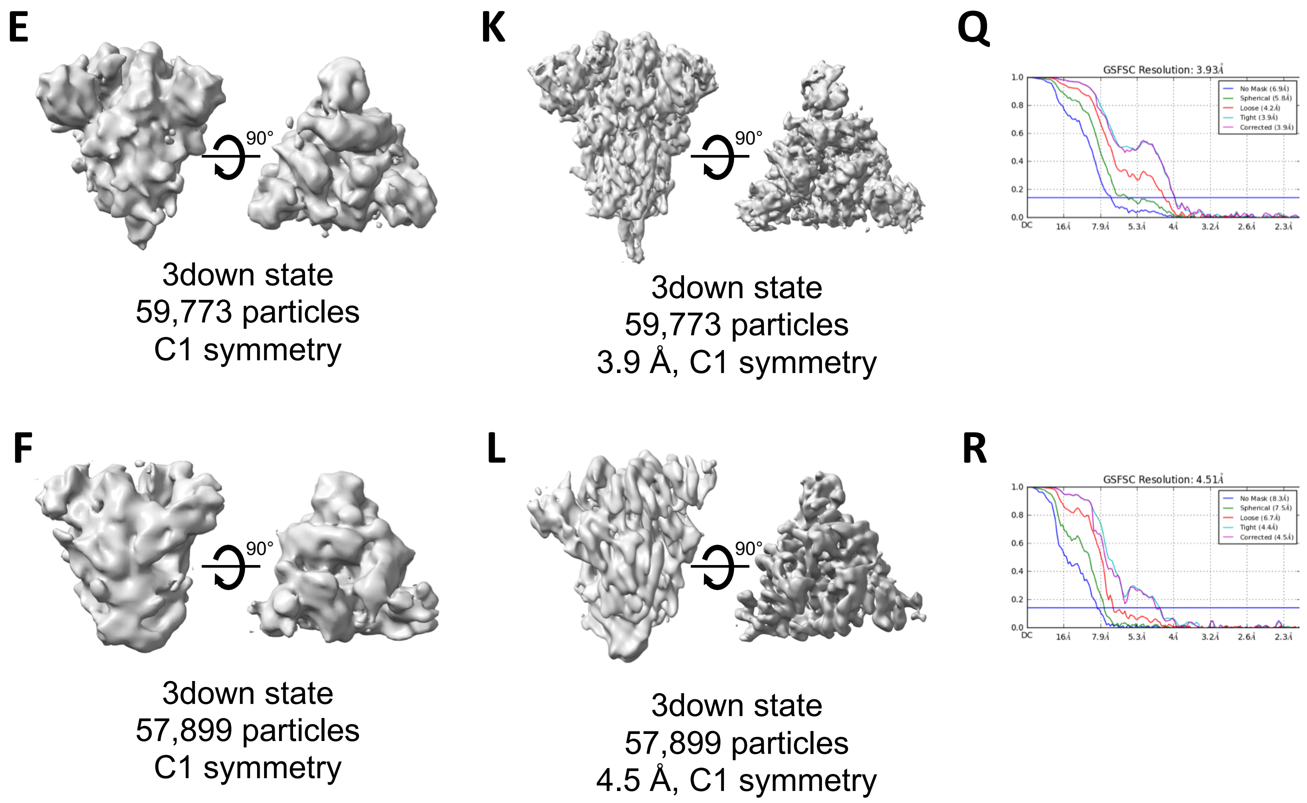


**Figure S7. Cryo-EM data processing details for S-GSAS/D614G 3-down subpopulations from Figure 3E. A-F.** *Ab initio* reconstructions for the 3-RBD-down subpopulations. **G-L.** Refined maps for the 3-RBD-down subpopulations. **M-R.** Fourier shell correlation curves for the 3-RBD-down subpopulations.

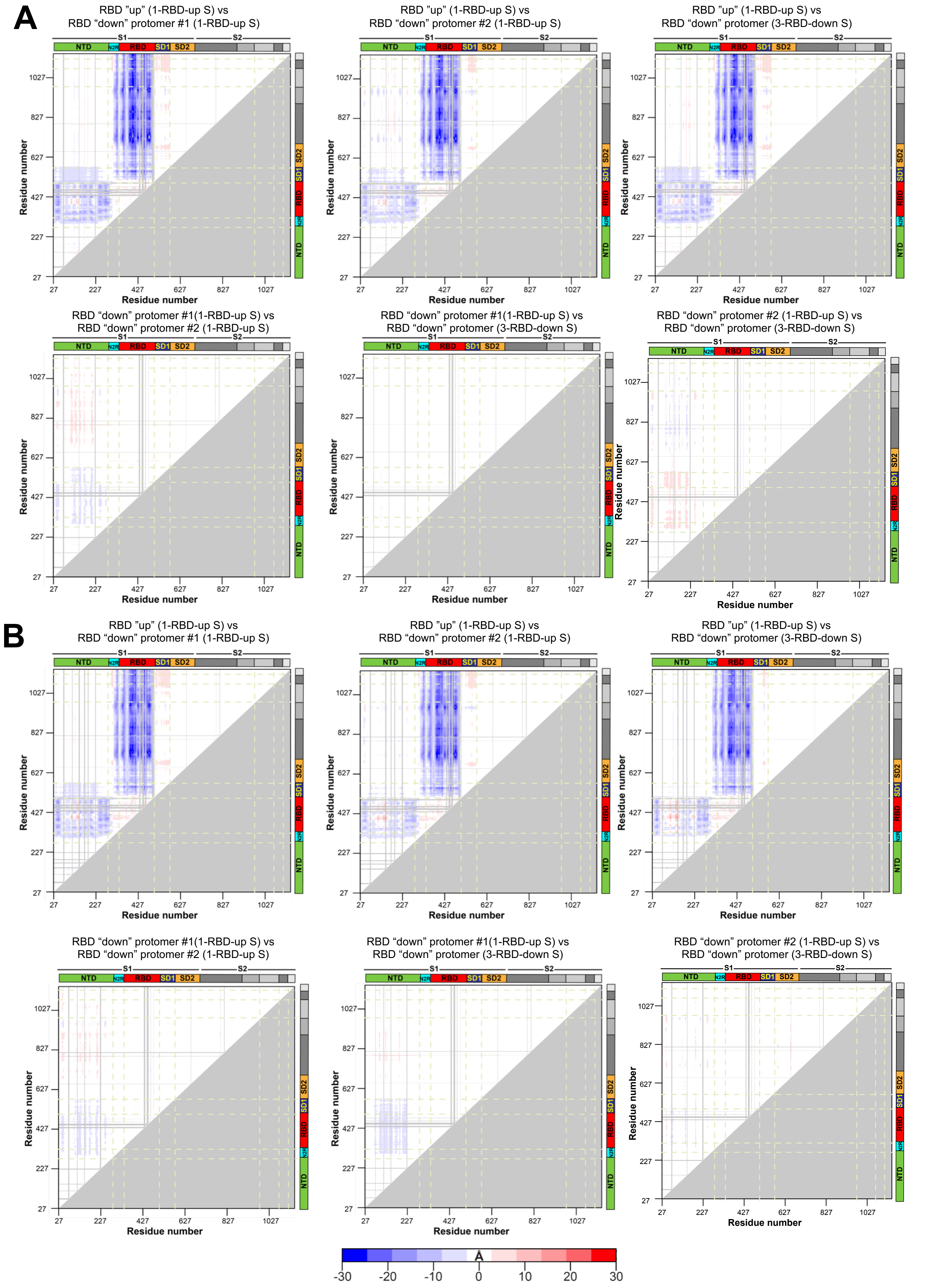

A
RBD "up" (1-RBD-up S) vs
RBD "down" protomer #1 (1-RBD-up S)
RBD "up" (1-RBD-up S) vs
RBD "down" protomer #2 (1-RBD-up S)
RBD "up" (1-RBD-up S) vs
RBD "down" protomer (3-RBD-down S)
RBD "down" protomer #1(1-RBD-up S) vs
RBD "down" protomer #2 (1-RBD-up S)
RBD "down" protomer #1(1-RBD-up S) vs
RBD "down" protomer (3-RBD-down S)
RBD "down" protomer #2 (1-RBD-up S) vs
RBD "down" protomer (3-RBD-down S)
B
RBD "up" (1-RBD-up S) vs
RBD "down" protomer #1 (1-RBD-up S)
RBD "up" (1-RBD-up S) vs
RBD "down" protomer #2 (1-RBD-up S)
RBD "up" (1-RBD-up S) vs
RBD "down" protomer (3-RBD-down S)
RBD "down" protomer #1(1-RBD-up S) vs
RBD "down" protomer #2 (1-RBD-up S)
RBD "down" protomer #1(1-RBD-up S) vs
RBD "down" protomer (3-RBD-down S)
RBD "down" protomer #2 (1-RBD-up S) vs
RBD "down" protomer (3-RBD-down S)
S1
S2
NTD
N2R
RBD
SD1
SD2
Residue number
27
227
427
627
827
1027
Å
-30
-20
-10
0
10
20
30

**Figure S8. Difference distance matrices (DDM) analysis of S-GSAS and S-RRAR/D614G fully cleaved by furin showing structural changes between different protomers.** The S-GSAS DDM are shown in **A** and S-RRAR/D614G in **B**. The blue to white to red coloring scheme is illustrated at the bottom.

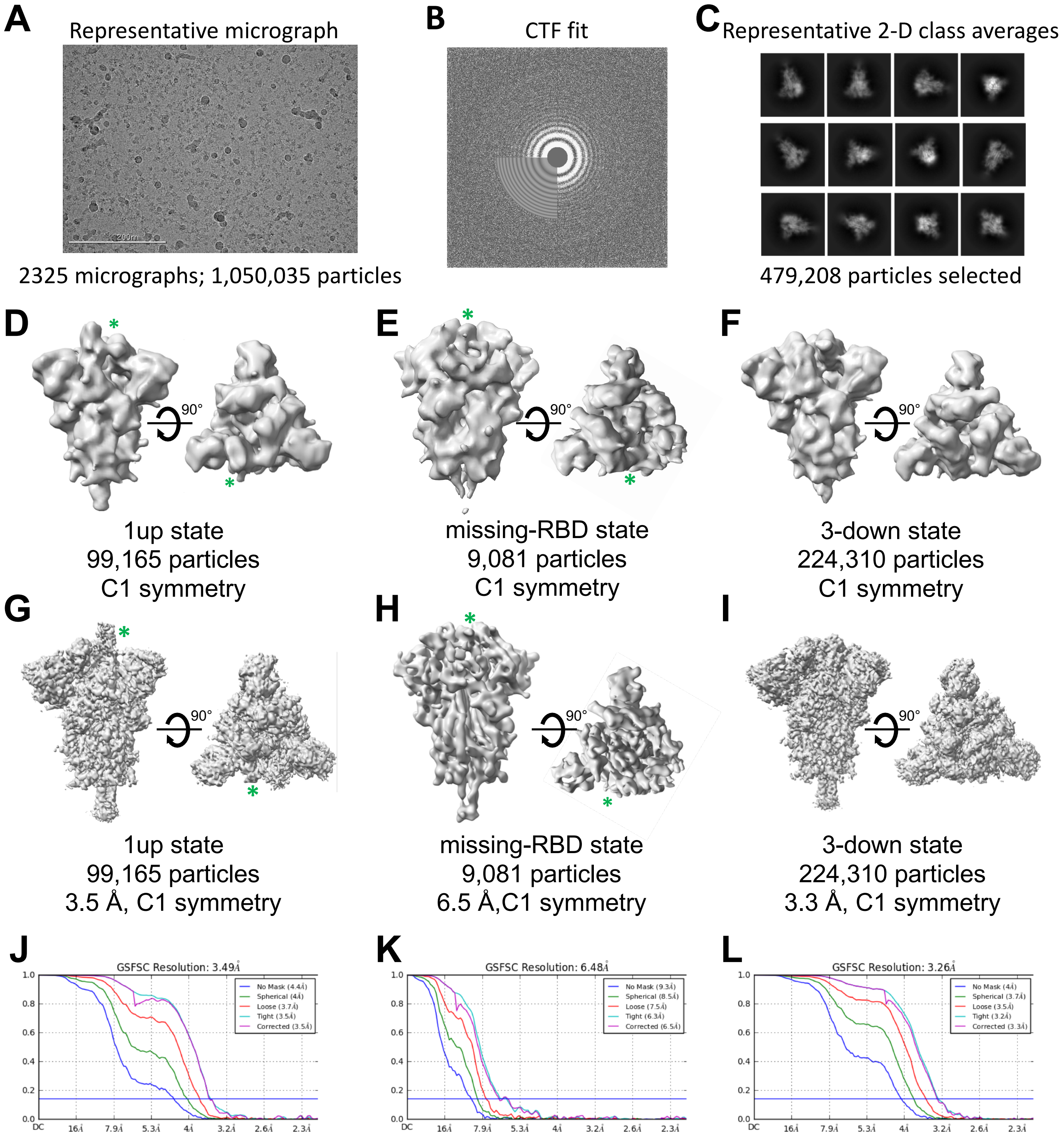


**Figure S9. Cryo-EM data processing details for S-RRAR/D614G ectodomain fully digested by furin. A.** Representative micrograph. **B.** CTF fit. **C.** Representative 2D class averages. **D-F.** *Ab initio* reconstructions for the **D.** 1-RBD-up state, **E.** missing-RBD state (the RBD in the up position or missing is identified by an asterisk) and **F.** 3-RBD-down state. **G-I.** Refined maps for the **G.** 1-RBD-up state, **H.** missing-RBD state (the RBD in the up position or missing is identified by an asterisk) and **I.** 3-down state. **J-L.** Fourier shell correlation curves for the **J.** 1-up state, **K.** missing-RBD state and **I.** 3- RBD-down state.

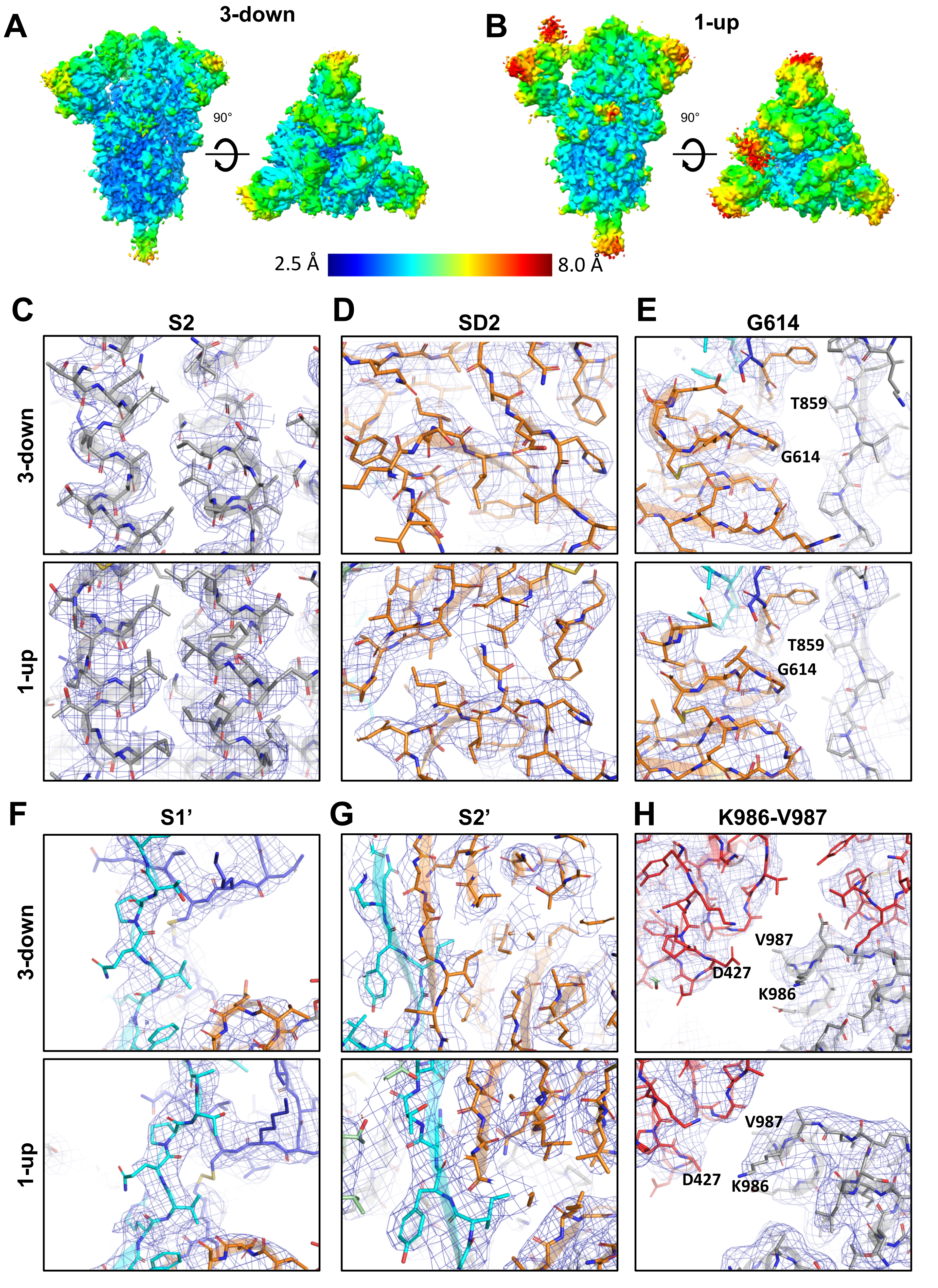

A
3-down
B
1-up
90°
90°
2.5 Å
8.0 Å
C
S2
D
SD2
E
G614
3-down
1-up
T859
G614
T859
G614
F
S1'
G
S2'
H
K986-V987
3-down
1-up
V987
D427
K986
V987
D427
K986

**Figure S10. Local map resolutions for the furin-cleaved S-RRAR/D614G ectodomain. A-B.** Refined cryo-EM maps colored by local resolution for the **A.** 3-RBD-down and **B.** 1-RDB-up states. **C-H.** Zoomed-in images showing representative views of the **C.** S2 subunit, **D.** SD2 subdomain, **E.** region around the D614 mutation, **F.** S1' "super" subdomain **G.** S2' "super" subdomain and **H.** K986-V987 regions in the 3-RBD-down (top) and 1-RDB-up spike structures (bottom). The cryo-EM map is shown as a blue mesh and the fitted model is in cartoon representation, with residues shown as sticks.
